# Transcriptome and translatome changes in germinated pollen under heat stress uncover roles of transporter genes involved in pollen tube growth

**DOI:** 10.1101/2020.05.29.122937

**Authors:** Laetitia Poidevin, Javier Forment, Dilek Unal, Alejandro Ferrando

## Abstract

Plant reproduction is one key biological process very sensitive to heat stress and, as a consequence, enhanced global warming poses serious threats to food security worldwide. In this work we have used a high-resolution ribosome profiling technology to study how heat affects both the transcriptome and the translatome of *Arabidopsis thaliana* pollen germinated *in vitro*. Overall, a high correlation between transcriptional and translational responses to high temperature was found, but specific regulations at the translational level were also present. We show that *bona fide* heat shock genes are induced by high temperature indicating that *in vitro* germinated pollen is a suitable system to understand the molecular basis of heat responses. Concurrently heat induced significant down-regulation of key membrane transporters required for pollen tube growth, thus uncovering heat-sensitive targets. We also found that a large subset of the heat-repressed transporters is specifically up-regulated, in a coordinated manner, with canonical heat-shock genes in pollen tubes grown *in vitro* and *semi in vivo*, based on published transcriptomes from *Arabidopsis thaliana*. Ribosome footprints were also detected in gene sequences annotated as non-coding, highlighting the potential for novel translatable genes and translational dynamics.

## INTRODUCTION

The evidence of a rapid global warming since the beginning of the industrial epoch is very solid (1) and its global impact seems to have anthropogenic origin, as paleoclimate reconstructions found no evidence of worldwide coherent warm or cold periods in the preindustrial times (2). According to the known drivers of climate change, global warming predictions have been made with pessimistic scenarios especially for agriculture in low latitudes (3). The negative impact of enhanced global warming on agriculture poses serious risks for future food security and demands smart food system management strategies (4). Under this scenario understanding how increased temperature impacts plant biology is an important task.

Plant reproduction is considered the most sensitive stage of plant development to the effects of global warming (5) and, for many crop plants, pollen seems to be particularly vulnerable to high temperatures (6–9). Pollen sensitivity to heat occurs not only throughout its development prior to anther dehiscence but also during germination and pollen tube growth (10,11). Upon landing on the stigmatic tissue of the pistil, pollen grains of flowering plants hydrate and germinate to immediately initiate pollen tube growth over long distances to deliver the male gametes to the egg and central cells in the embryo sac for the double fertilization process (12). Growth of the pollen tube from the stigma to the ovules occurs at extremely rapid rates and requires a fine regulation of subcellular processes such as: well-ordered vesicle transport, precise cytoskeleton arrangements, strict and local pH establishment, proper energy management with amyloplast and mitochondrial logistics, and delicate regulation of regulatory ion fluxes for osmotic adjustment (K^+^ transport) and signaling (Ca^2+^ transport). This impressive machinery demands exquisite coordination among all these processes for the rapid transport of germinal sperm cells to the micropyle (13,14). One important aspect of this optimized cellular growth is the presence of local pH gradients with an acidic apex region and an alkaline band in the clear zone followed by a subapical region of acidic pH (15,16). This complex organization of events demands an optimized protein synthesis machinery (17), as critical protein components like calreticulin have been shown to be continuously synthesized during pollen tube growth (18).

Ultrastructural analysis of pollen tube growth at high temperature revealed alterations in rough-endoplasmic reticulum, Golgi apparatus and mitochondria (19) thus supporting the notion that protein translation, vesicle transport and energy depletion during pollen tube growth may be limiting steps in the adaptation to high temperature conditions. Molecular studies have uncovered heat responses in pollen subjected to high temperature similar to other plant cell types including the heat shock response, the unfolded ER protein response or the formation of reactive oxygen species (20). In addition, a number of untargeted transcriptomics, metabolomics, and proteomics studies have been performed to characterize the response of pollen under heat stress conditions at the molecular level (21–24). However, the limitations of the proteomics and metabolomics technologies compared to high-resolution approaches based on next generation sequencing of nucleic acids (NGS) techniques (25), qualify these NGS approaches as very appropriate to gain deeper insight in the response of pollen to heat stress conditions.

A breakthrough in the studies of mRNA translation was achieved with the development of the ribosome profiling technique (also called Ribosome Footprinting, Riboprofiling or Ribo-Seq), a ribosome-centric approach to isolate and sequence ribosome-protected mRNA fragments (RPFs) after RNase treatment that provides a quantitative profile of translation across the transcriptome with sub-codon resolution (26). The advantage of Riboprofiling is that, in parallel with massive sequencing of ribosome footprints, total RNA is also subjected to high-throughput sequencing thus allowing the direct comparison of the transcriptome, the translatome and the calculations of translational efficiencies for the complete set of transcripts (27).

Here we have used Riboprofiling with Arabidopsis pollen germinated *in vitro* under both optimum and high temperature conditions. We provide high-resolution transcriptome and translatome profiles of pollen tubes germinated at basal temperature, and show their changes in response to heat stress. We have found that *in vitro* germinated pollen responds to high temperature by up-regulating heat shock genes in a similar manner to the vegetative organs. Heat stress also down-regulated specifically many membrane transporters, including K^+^ and carbohydrate co-transporters, thus finding a suitable explanation to the deleterious effects of high temperature on pollination. A subset of these transporters is up-regulated during *semi in vivo* pollen development, when heat shock genes are also expressed, illustrating the importance of these heat-sensitive functions during plant fertilization. Moreover, we have found specific regulations at the translational level and the unexpected presence of ribosome footprints on non-coding RNAs. Thus, our analysis of combined transcriptome and translatome is revealing novel insights on pollen responses to heat stress, including a tight connection between heat shock gene expression and membrane transporters that promote pollen tube growth and male fertility.

## MATERIAL AND METHODS

### Large-scale pollen collection using vacuum filtration

About 20.000 *A. thaliana* (Col-0) plants were grown in greenhouse for 4 weeks in a mixture of 25% vermiculite, 25% perlite and 50% soil under long-day photoperiod cycles. Pollen collection was performed by vacuum filtration following described protocols (28). Pollen was then scrapped off the 6μm nylon mesh, and kept to dry overnight under a chemical hood in a 2mL tube. After weighing, a silica bead was added to the pollen tube and the samples were placed at −20°C. Harvests were done every second day for about 2 weeks. Yields ranged from 10 to 55 mg of pollen per day. A small aliquot from each sample was kept in a second tube for pollen viability and contamination assays. The whole experiment was repeated 3 times to obtain 3 biological replicates.

### Pollen growth assays

The pollen tube growth was assayed using published protocols (29). Frozen pollen grains were first rehydrated for 1h in a humid chamber after removing the silica beads. Pollen was then germinated on an agar medium containing 10% sucrose, 0.01% boric acid, 1 mM MgSO_4_, 5 mM CaCl_2_, 5 mM KCl, and 1.5 % low-melting agarose, pH 7.5 (adjusted with 0.1M NaOH). For microscopic pollen examination, 1 mL of melted germination medium was spread on a glass microscope slide to build a flat pad where pollen can be spread after the agarose is solidified. The slides were placed inside a moisture incubation chamber to avoid dehydration of the medium, and incubated in the dark at 24°C or 35°C, for the indicated time. Pictures were taken with a Leica DM5000 microscope. Quantifications of pollen germination and tube length were performed with the IMAGEJ Software (30) using the cell counter and NeuronJ plug-ins (31). Germinated pollen was scored positively for grains with a pollen tube length of at least the pollen diameter. For the Riboprofiling experiments, the protocol was scaled up to 10 cm-square petri dishes containing 30 mL of pollen germination medium. 25 mg of pollen grains were spread per plate with a soft brush in a moisture incubation chamber, sealed, and placed in the dark at 24°C or 35°C for 5 hours before being collected. 3 plates per temperature were prepared from each of the biological replicates.

### Preparation of RNA and RPF libraries for Riboprofiling

Pollen was scrapped from the plates and ground in liquid nitrogen in an optimized ice-cold extraction buffer for Riboprofiling experiments (32) containing: 0.1% Tris-HCl pH8, 1% sodium deoxycholate, 40 mM KCl, 20 mM MgCl_2_, 10% polyoxyethylene-10-tridecyl ether, 1mM DTT,100 μg/mL cycloheximide, 10U/mL DNAse I (Illumina, USA). After centrifugation, the supernatants were transferred into a pre-chilled tube and split into 100 μL (for RNA library) and 200 μL (for RPF library) aliquots. Estimation of RNA concentration was performed with NanoDrop ND1000 (Thermo Fisher Scientific, USA). The libraries were obtained using the Illumina® TruSeq® Ribo Profile Mammalian kit (Illumina), with slight modifications to the standard protocol. The 100 μL samples of total RNA were denatured by adding SDS to 1% final concentration and kept aside until needed. In parallel, generation of ribosome protected fragment consisted of digesting 200 μg of RNA with 100 U of TruSeq Ribo Profile Nuclease for 1h at 21°C, with shaking at 300 rpm. Reaction was stopped by adding 1U/μL of SUPERase In RNase Inhibitor (Thermo Fisher Scientific, USA), then ribosomes were purified by size exclusion columns using MicroSpin S-400 columns following provider’s recommendations (GE Healthcare, UK). Recovered ribosomes were denatured by adding SDS to 1% final concentration, and then both ribosome-bound RNA and denatured total RNA samples were purified with RNA Clean & concentrator-25 kit (Zymo Research, USA). RPFs were size-fractionated on 15% urea polyacrylamide gel electrophoresis, by recovering fragments of 28 to 30 nt after gel staining with SYBR Gold (Thermo Fisher Scientific, USA), using Truseq Ribo Profile RNA Control as reference. Purified RPFs and total RNAs were rRNA depleted using the Ribo-Zero Magnetic bead kit (Illumina, USA) by adding 50 μL or 100 μL of magnetic beads per RPF or RNA samples respectively, according to manufacturer’s instructions. Heat fragmentation of total RNA, end repair of RNA and RPF as well as downstream steps for 3’ adapter ligation, adaptor removal, reverse transcription, PAGE purification of cDNA, and cDNA circularization were done following strictly the Illumina kit protocol. The cDNA Libraries were amplified by 13 and 15 PCR cycles for RNA and RPF, respectively. Library fragments of expected size were purified by non-denaturing polyacrylamide gel electrophoresis and, after purification, DNA integrity and concentration were checked using Bioanalyzer 2100 expert High Sensitivity DNA Assay (Agilent Technologies, Inc). Equimolar (5nM) pools of libraries were prepared for 100 bp paired-end sequencing on Illumina HiSeq4000 platform at Macrogen (Korea).

### Bioinformatic data analysis

To determine which of the paired files corresponded to reads from the coding strand, htseq-count v0.10 was used and the files were kept for downstream analysis. After quality analysis with FastQC v0.11.5, reads were processed for adaptor and low quality regions removal with cutadapt v1.16 (33). Clean, trimmed reads ranging between 20-40 nt for RNA samples and 20-30 nt for RPF samples were selected for downstream analysis. Reads corresponding to rRNA, tRNA, snRNA, and snoRNA sequences were identified and removed with bowtie2 v2.3.2 (34) using the set of rRNA, tRNA, snRNA, and snoRNA TAIR10 sequences downloaded from Ensembl Plants (35). The remaining reads were mapped to the TAIR10 Arabidopsis reference genome sequence using HISAT2 v2.1.0 (36), and entries for reads mapping to more than one locus were excluded with samtools v1.5 (37). For RPF samples, only 27-28 nt uniquely mapped reads were selected for downstream analysis using the Linux command line ‘awk’. The final sets of RNA 20-40 nt and RPF 27-28 nt uniquely mapped reads were assigned to specific genes, 5’UTRs, CDSs, and 3’UTRs using htseq-count v0.10 (38) and the last Arabidopsis TAIR10 genome annotation from Araport11 (39) containing 37,336 genes. Correlation heat-maps were obtained with multiBamSummary and plotCorrelation, from the deepTools v3.1.0 package (40). PCA analysis was performed in R v3.4.4, using the cluster v2.1.0 (41), Biobase v2.38.0 (42), qvalue v2.10.0 (43), and fastcluster v1.1.25 (44) packages. Ribowave v1.0 (45) was used for the study of periodicity of RPF reads, and Xtail v1.1.5 (46) for the differential transcriptional/translational analysis and also for the studies of differential translational efficiency. The fold change comparisons always refer to the 35°C data (treatment) versus the 24°C data (no treatment). The GO term enrichment analysis was performed using the web-based g:Profiler toolset (47) with the g:SCS threshold method and either annotated genes or custom list of genes when indicated. RStudio v1.2.5001 (48) was used with the ggplot2 library for correlation plots and periodicity graphs, and with the UpSetR library for the intersection set analysis. To obtain the heat-map viewer for gene expression under different pollen developmental stages we used the web-based tool ‘Arabidopsis Heat Tree Viewer’ (http://arabidopsis-heat-tree.org/). Comparison of gene lists with Venn diagrams was performed with the web tool Venny 2.1 (49). Graphical visualization of the read coverage on mapped genes was performed with the Integrative Genomics Viewer (IGV) (50).

## RESULTS

### Establishing the protocol for Riboprofiling studies of pollen response to elevated temperature

We first established a protocol for large-scale pollen collection and *in vitro* pollen germination necessary to perform the Riboprofiling studies. We grew around 60,000 *A.thaliana* Col-0 plants under greenhouse conditions to collect pollen by vacuum filtration with a modified vacuum cleaner (Figure 1A, left) as previously published (28). Pollen was collected and, after dehydration, stored in dry aliquots at −20°C (Figure 1A, middle and right panels) to allow the synchronous processing of the samples. A small amount of every frozen aliquot, after proper rehydration, was tested for germination under *in vitro* conditions. Only non-contaminated samples showing good germination percentages were used to quantify germination and pollen tube growth under optimum temperature (24°C) and limiting high temperature conditions (35°C). As shown in Figure 1B (left), maximum germination was achieved after 5 hours of incubation at both temperatures, and germination was negatively affected by the elevated temperature. Pollen tube growth showed a severe growth inhibition at 35°C from early growth stages (Figure 1B, right). We therefore selected a time point of 5 hours of incubation to collect 3 biological replicates of germinated pollen both at 24°C and 35°C for Riboprofiling analysis. Once *in vitro* germinated pollen was collected from the plates, every replicate was divided in two samples for parallel isolation of total RNA (RNAs) and ribosome protected fragments (RPFs), therefore a total of 12 libraries were prepared for next-generation sequencing (Figure 1C).

**Figure 1.**
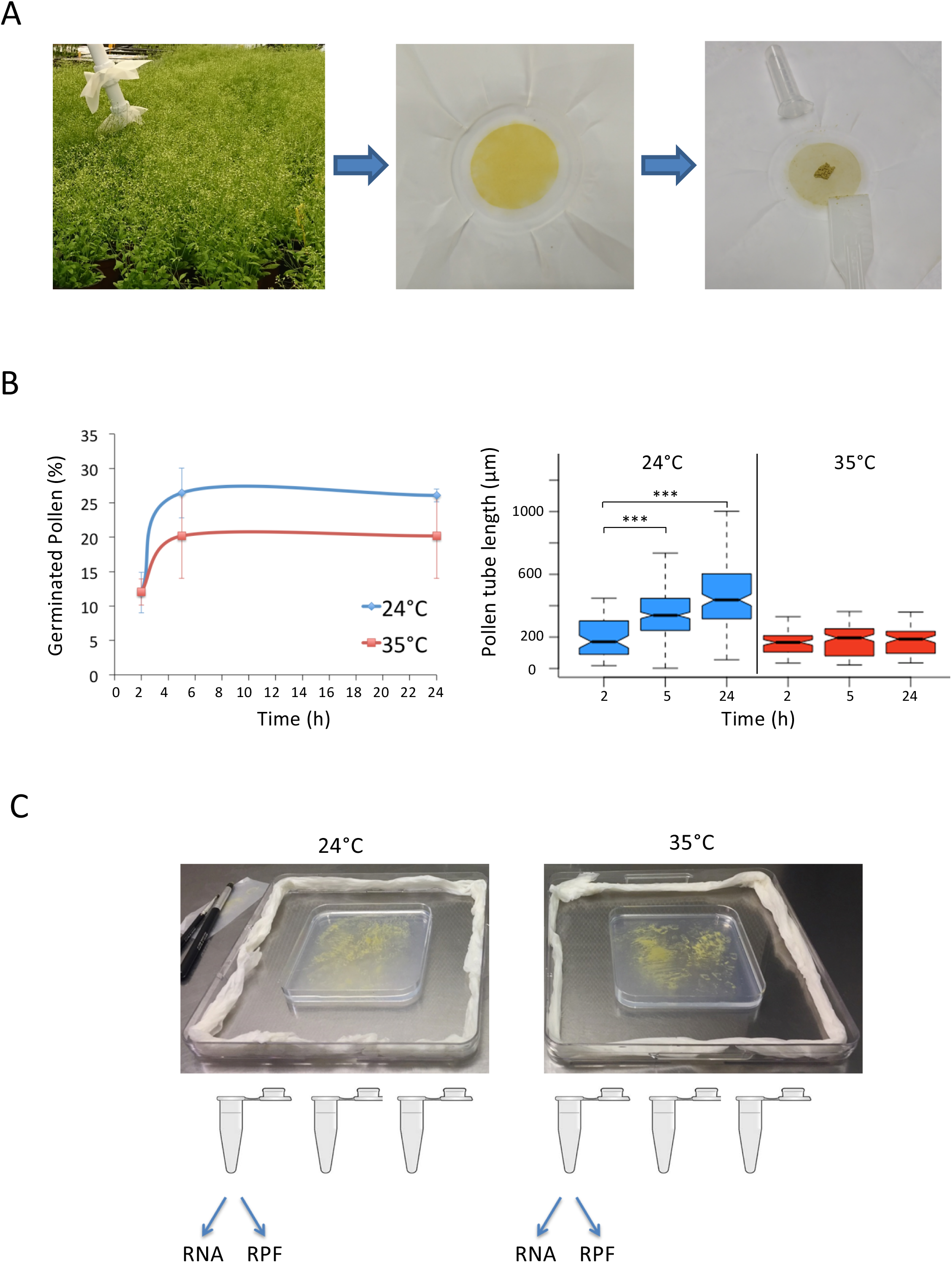
Protocol for Riboprofiling of germinated pollen at different temperatures. A, large scale growth of *Arabidopsis thaliana* plants under greenhouse conditions and pollen collection by vacuum filtration. B, time course of germination rate and pollen tube growth were scored for at least 250 and 75 pollen grains respectively. Error bars represent standard deviation of 3 independent experiments, and asterisks indicate significant differences (p < 0.001) with respect to time 2 hours after *t*-test analysis of the same biological replicates. C, rehydrated pollen was germinated and grown in humid chambers at two different temperatures for the simultaneous isolation of total RNA and ribosome-protected RNA fragments (RPF) from 3 biological replicates.

### Riboprofiling shows enrichment of coding sequences in germinated pollen

High-throughput sequencing was performed using Illumina HiSeq4000 yielding a total of over 3100 million paired-end 100 base reads for the 12 libraries. The numbers of reads for each library after several bioinformatics processing steps are summarized in Table 1. The correlation and PCA analyses among the different RNA and RPF libraries are shown in Supplementary Figure S1. The data show very good cluster separations between the type of library (RNA and RPF) and between the temperature treatments (24°C and 35°C), and the libraries displayed very high correlation coefficients with values above 0.95 among the 3 replicates in all cases. We then analysed the distribution of read lengths for the RNA and the RPF reads shown in Figure 2A. Whereas RNA reads are spread in a large range size from 20 to 40 nucleotides, the RPF reads are restricted to shorter lengths showing a notable accumulation around 27 and 28 nucleotides as expected for ribosome footprint protection in *A.thaliana* (32).

**Figure 2.**
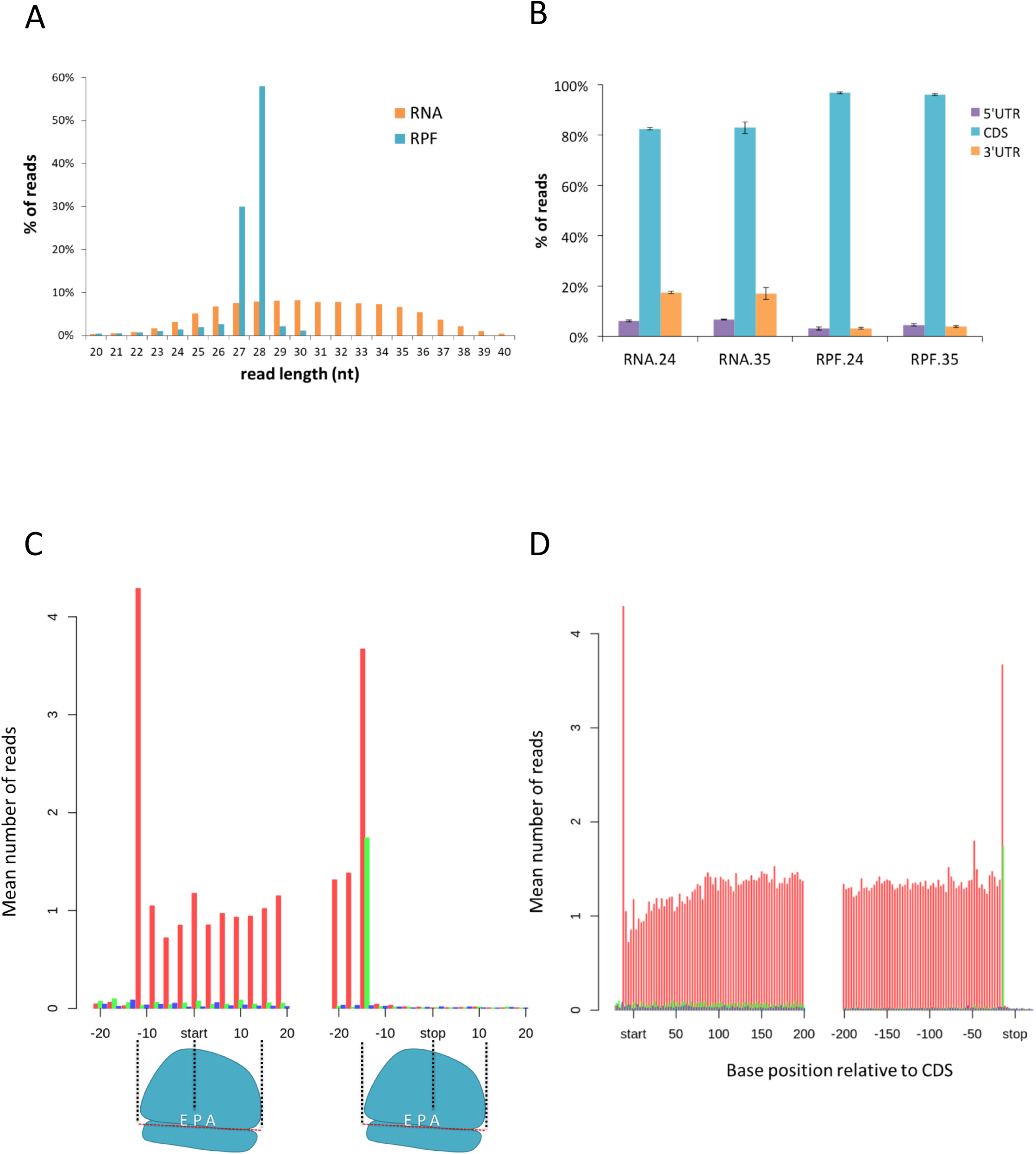
Features of the RNA and RPF libraries of germinated pollen. A, size distributions of the RNA and RPF reads for all the sequenced libraries combined. B, distribution of the reads in genomic features for all the libraries comparing coding regions (CDS) and untranslated regions (5’UTR and 3’UTR). C, meta-gene coverage values of the 28-nt reads of the RPF24.1 library showing the inferred position of the initiating and terminating ribosomes. D, the same values are shown for the whole coding region. The main frame according to the start position is shown in red, and the other two frames are shown in green and blue.

**Table 1.**
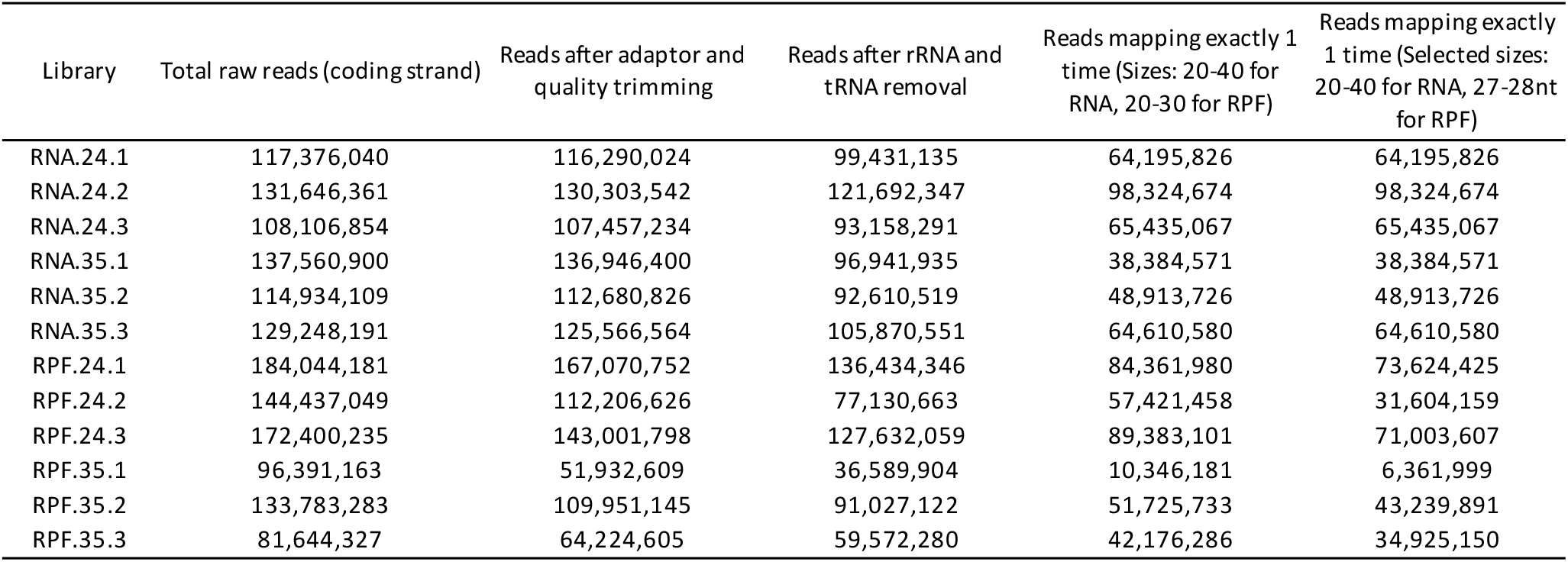
Number of reads per sequenced library remaining after bioinformatics processing. NGS library statistics. Data represent the remaining reads for each massively-sequenced library after the required bioinformatics steps of cleaning, mapping and size-selection.

| Library | Total raw reads (coding strand) | Reads after adaptor and quality trimming | Reads after rRNA and tRNA removal | Reads mapping exactly 1 time (Sizes: 20-40 for RNA, 20-30 for RPF) | Reads mapping exactly 1 time (Selected sizes: 20-40 for RNA, 27-28nt for RPF) |
| --- | --- | --- | --- | --- | --- |
| RNA.24.1 | 117,376,040 | 116,290,024 | 99,431,135 | 64,195,826 | 64,195,826 |
| RNA.24.2 | 131,646,361 | 130,303,542 | 121,692,347 | 98,324,674 | 98,324,674 |
| RNA.24.3 | 108,106,854 | 107,457,234 | 93,158,291 | 65,435,067 | 65,435,067 |
| RNA.35.1 | 137,560,900 | 136,946,400 | 96,941,935 | 38,384,571 | 38,384,571 |
| RNA.35.2 | 114,934,109 | 112,680,826 | 92,610,519 | 48,913,726 | 48,913,726 |
| RNA.35.3 | 129,248,191 | 125,566,564 | 105,870,551 | 64,610,580 | 64,610,580 |
| RPF.24.1 | 184,044,181 | 167,070,752 | 136,434,346 | 84,361,980 | 73,624,425 |
| RPF.24.2 | 144,437,049 | 112,206,626 | 77,130,663 | 57,421,458 | 31,604,159 |
| RPF.24.3 | 172,400,235 | 143,001,798 | 127,632,059 | 89,383,101 | 71,003,607 |
| RPF.35.1 | 96,391,163 | 51,932,609 | 36,589,904 | 10,346,181 | 6,361,999 |
| RPF.35.2 | 133,783,283 | 109,951,145 | 91,027,122 | 51,725,733 | 43,239,891 |
| RPF.35.3 | 81,644,327 | 64,224,605 | 59,572,280 | 42,176,286 | 34,925,150 |

After mapping to the *A.thaliana* reference genome, the RNA and RPF read distributions were computed on 5’UTR, coding regions and 3’UTR, showing that ribosome footprints map to higher extent to coding sequences and to lesser extent to untranslated regions compared to RNA reads (Figure 2B). The bias of RPFs towards the translated frame can be used to infer the position of the ribosomal peptidyl-site (P-site) as a very good proxy to determine whether the ribosome is in frame with either start or stop codons. As an example, the meta-gene computing analysis of RPF distribution for 28 nucleotide fragments of one library is shown in Figures 2C and D. The data clearly show a well-defined trinucleotide (3-nt) periodicity with the enhanced peaks corresponding to the start and stop sites, and depict the expected coverage of −12 and −15 nucleotide position for the initiation P-site and termination A-site respectively, as it has been shown for high quality *A.thaliana* RPF libraries (32). The data corresponding to the complete set of RPF libraries and size lengths ranging from 26 to 30 nucleotides is shown in Supplementary Figure S2. We found a very strong periodicity for the fragments of 27 and 28 nucleotides in all the libraries therefore we selected those reads for further analysis.

### The transcriptome and translatome landscapes of germinated pollen at optimum temperature

The gene-specific read counts for RNA and RPF were normalized using the *transcript per kilobase million* (TPM), as it shows better consistency than *reads per million per kilobase* (RPKM) normalization for comparisons among different samples (51,52). The complete set of raw counts and normalized TPM data for the *A.thaliana* genome have been deposited at GEO with the accession number GSE145795. Although a minimum value of 1 TPM has been established as a threshold to qualify a gene as expressed in other plant tissues (53) and metazoans (54), we chose the more conservative value of 2 TPM. We then defined the *A.thaliana* pollen transcriptome at 24°C as the set of genes having at least 2 TPM in the average of the 3 replicates of RNA samples at 24°C, with the condition that at least 2 of the 3 replicates should also score a minimum of 2 TPM values. The table of 6098 genes corresponding to the germinated *A.thaliana* pollen transcriptome at 24°C can be found as Supplementary Table S1_Tab1.

With the aim to validate our transcriptome list of germinated pollen at 24°C, we first checked the expected absence of photosynthetic genes in our samples of heterotrophic tissue by analysing the expression values of 30 light-harvesting complex genes of photosystem I and II (55) and we found all of them either not detectable or with TPM values close to zero. Next, we compared the transcriptome gene list with previously reported Arabidopsis pollen gene expression at different developmental stages (56–62). To find the common set of pollen transcriptome genes among the different gene lists, we used the UpSetR bioinformatics tool to visualize the intersecting sets (63). Figure 3A shows the number of genes for all possible intersections among the different lists, with 2825 genes common to all of them. In addition to the set of genes common to all datasets, our defined pollen transcriptome gene list also includes genes catalogued as pollen-expressed genes in the different published sets. The differences very likely account for variations among the pollen developmental stage, growth conditions and RNA extraction procedures, but also for the threshold values used to define a gene as being expressed.

**Figure 3.**
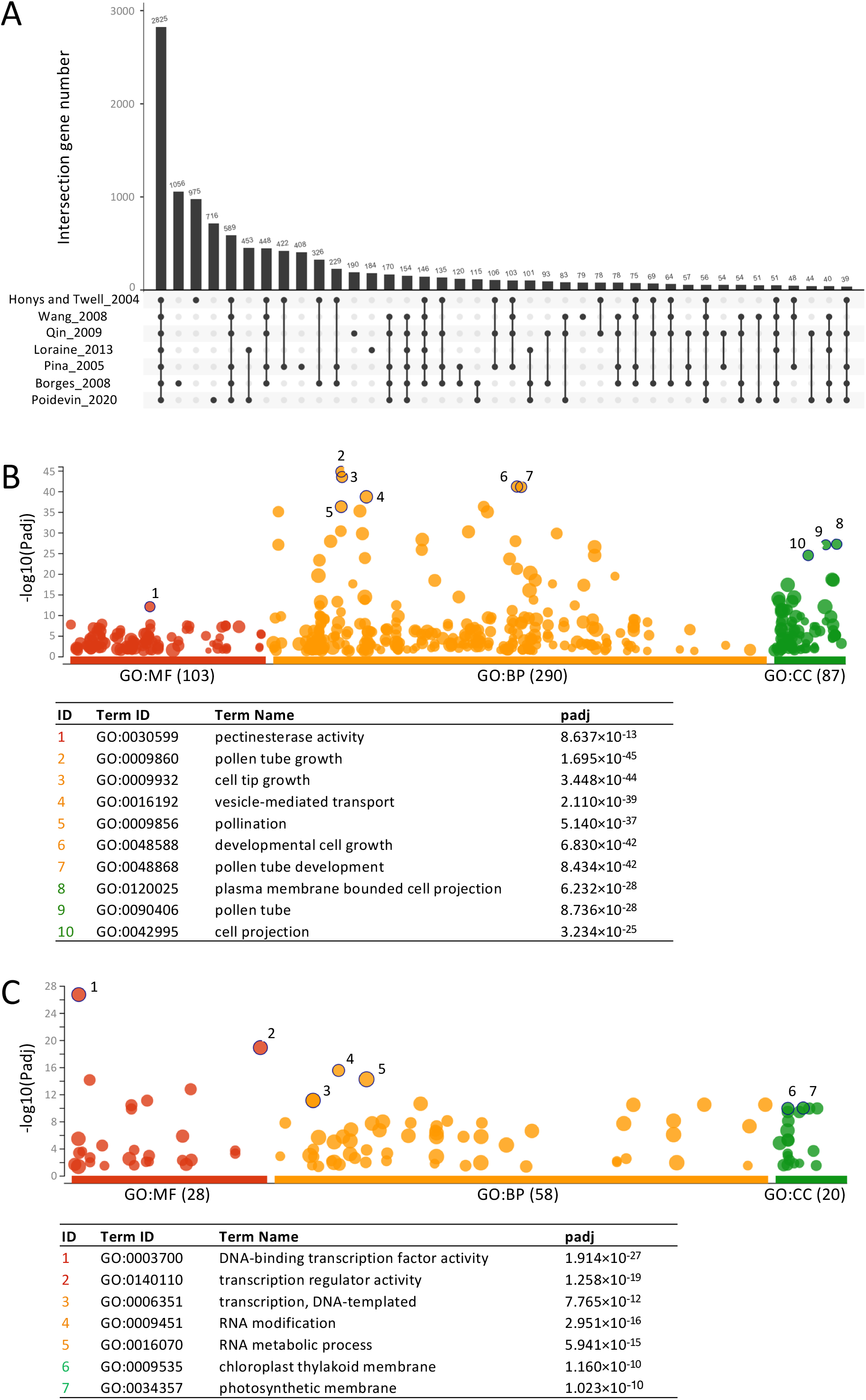
Characterization of the germinated pollen transcriptome at optimum temperature. A, the UpSetR tool was used to identify the intersection set of genes (black dots) among previously published pollen transcriptomes compared with our own transcriptome gene list. Unique set of genes are indicated by unconnected dots. B, the Manhattan plot of the enrichment of GO terms in our transcriptome gene list was generated with the g:Profiler toolset highlighting the most significant cases in the different sources indicated by colours. MF refers to ‘Molecular Function’ in red, BP to ‘Biological Process’ in orange, and CC to ‘Cellular Component’ in green. The dot size is proportional to the number of genes included in the term. C, the g:Profiler plot was used the same way to identify underrepresented GO terms in our transcriptome gene list of germinated pollen at optimum temperature. The GO analysis were performed against the *Arabidopsis thaliana* genome in both cases.

To further check whether the set of genes properly define the pollen transcriptome, we used the web-based g:Profiler toolset (47) to find enriched and underrepresented biological categories in our transcriptome gene list. The advantage of this tool, compared to other methods, is that it is a fast, flexible, and well-updated service providing ready-to-use intuitive Manhattan plots. The results shown in Figure 3B, display the presence of expected overrepresented terms such as ‘pectinesterase activity’, ‘pollen tube growth’, ‘vesicle-mediated transport’ and ‘cell projection’ among others. On the contrary, Figure 3C shows underrepresented GO terms including ‘transcription regulator activity’, ‘RNA metabolic process’, ‘chloroplast thylakoid membrane’ and ‘photosynthetic membrane’, expected down-regulated terms during pollen development. However, as it is well-known, the GO term lists should be taken with caution in particular for large gene sets as for instance it has been shown that some pectinesterases are down-regulated during pollen tube growth and at the same time some transcription factors are up-regulated (60,64). In fact, it has been reported that inhibition of transcription blocks only partially pollen germination and pollen tube growth whereas translation is fully required for both processes (58,65).

Once we confirmed that our transcriptome list adequately reflected the mRNA levels of genes involved in pollen germination and growth, we wanted to compare these data with those provided by RPF libraries. These libraries represent the quantitative translational status of the gene set or the translatome, defined as the ribosome-bound mRNA fragments considered to be actively translated in a given situation. We defined the *A.thaliana* pollen translatome at 24°C as the set of genes having at least 2 TPM in the average of the 3 replicates of both RNA and RPF samples at 24°C, with the condition that at least 2 of the 3 replicates should also score a minimum of 2 TPM values. The translatome gene list for germinated pollen at 24°C (shown in the Supplementary Table S1_Tab2), accommodates a total of 4782 genes representing our best approximation to the proteome of germinated pollen. We were intrigued by the large list of 1316 genes absent in the translatome but present in the transcriptome, shown in Supplementary Table S1_Tab3. Therefore we performed a GO analysis with this gene list and identified several terms related to ‘DNA replication’, ‘DNA repair’ and ‘chromosome organization’, which are typical GO terms for sperm cells (57) as shown in Supplementary Figure S3 A. Supplementary Figure S3 B shows one example of strong translational repression with the IGV visualization of RNA and RPF reads for for the gene AT5G16020 encoding a gamete expressed protein GEX3. This indicates that a strict translational regulation occurs during pollen germination to avoid translation of unnecessary transcripts at this developmental stage.

The use of proteomic approaches has revealed only a modest correlation between mRNA transcript and protein levels (66–68), therefore we carried out a correlation plot analysis between the germinated pollen transcriptome and translatome (Figure 4A). The correlation R value of 0.88 suggests a rather high correspondence between the transcriptome and the translatome for germinated pollen at 24°C, although some exceptions can be detected. To visualize the differences in the translational profile revealed by the RPF values we used RiboWave (45), a pipeline able to denoise the original RPF signal and extract the 3-nt periodicity of the reads known as Periodic Footprint P-site or PF P-site. In Figure 4B we plotted the PF P-site values of two gene examples taken from Figure 4A with similar transcriptional value but obvious differences at the translational level, to show that the main differences are visualized by alterations in PF P-site values throughout the coding region. Altogether we have obtained representative gene lists of transcriptome and translatome of *A.thaliana* pollen germinated at optimum temperature (24°C).

**Figure 4.**
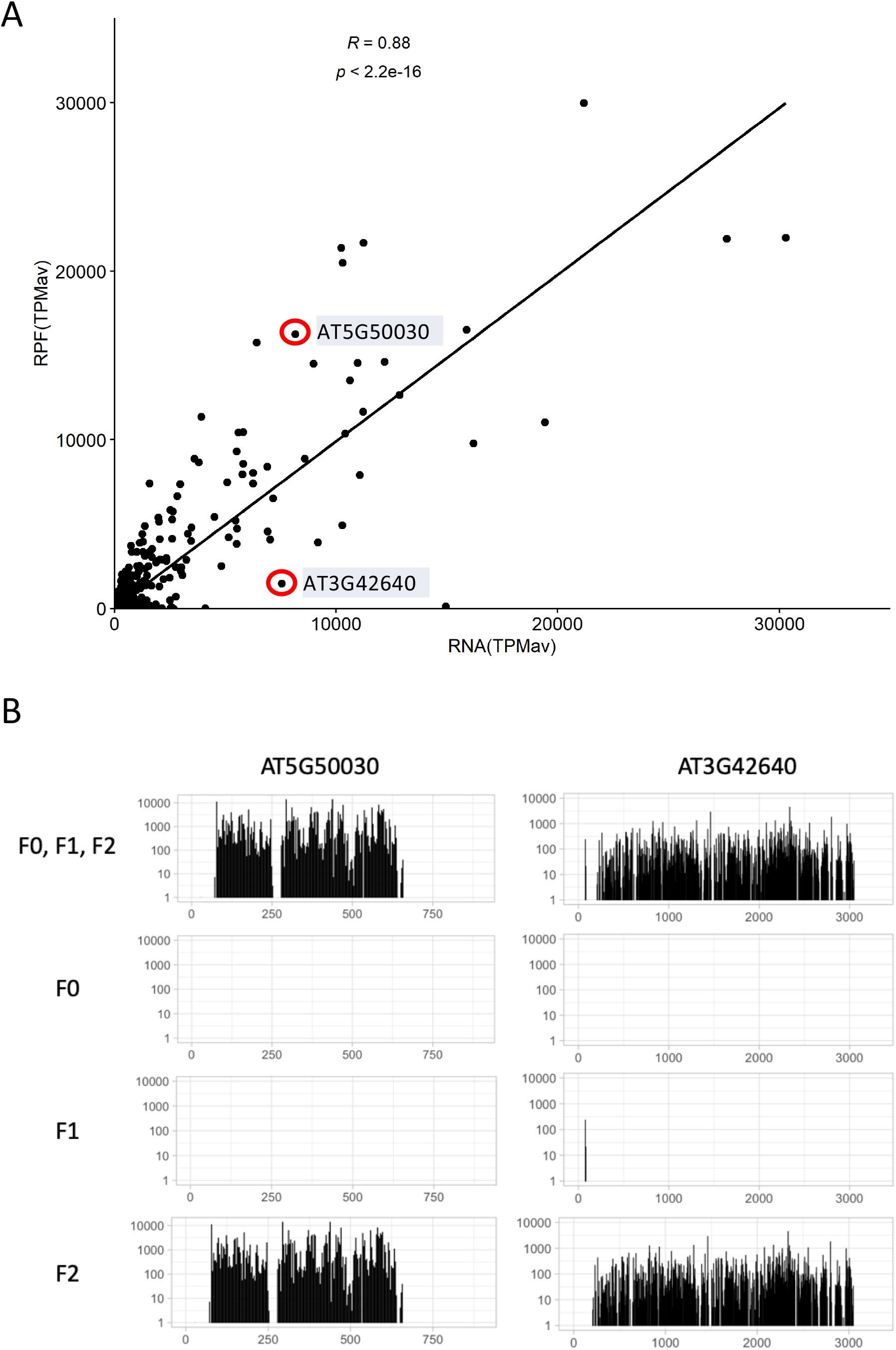
Characterization of the germinated pollen translatome at optimum temperature. Panel A shows the correlation plot of the germinated pollen translatome (RPF) versus the transcriptome (RNA) at optimum temperature, with the average TPM value of both sets of genes. Highlighted with red circles are two examples of genes with similar RNA but different RPF values. Panel B shows the periodic PF P-site distribution across both genes highlighted in Panel A in the 3 reading frames (F0, F1, F2) or for every reading frame separately.

### Transcriptional and translational alterations of germinated pollen by high temperature

The RNA and RPF libraries of germinated pollen at 35°C were computed the same way as it was done previously with the 24°C libraries, allowing us to generate both the transcriptome and translatome gene lists of germinated pollen at 35°C shown in Supplementary Table S2_Tab1 and Supplementary Table S2_Tab2, respectively. Similar to the libraries at optimum temperature, the gene list numbers at the restrictive temperature of 35°C were 6089 for the transcriptome and 4742 for the translatome. We compared the intersection sets among the 4 gene lists generated (transcriptome 24, translatome 24, transcriptome 35 and translatome 35) using Venn diagrams (Figure 5A). A total of 4386 genes were common to all 4 sets of genes; 1415 genes present in the transcriptomes were absent in the translatome lists, and 919 genes were exclusively expressed at either 24°C (464 genes) or 35°C (455 genes).

**Figure 5.**
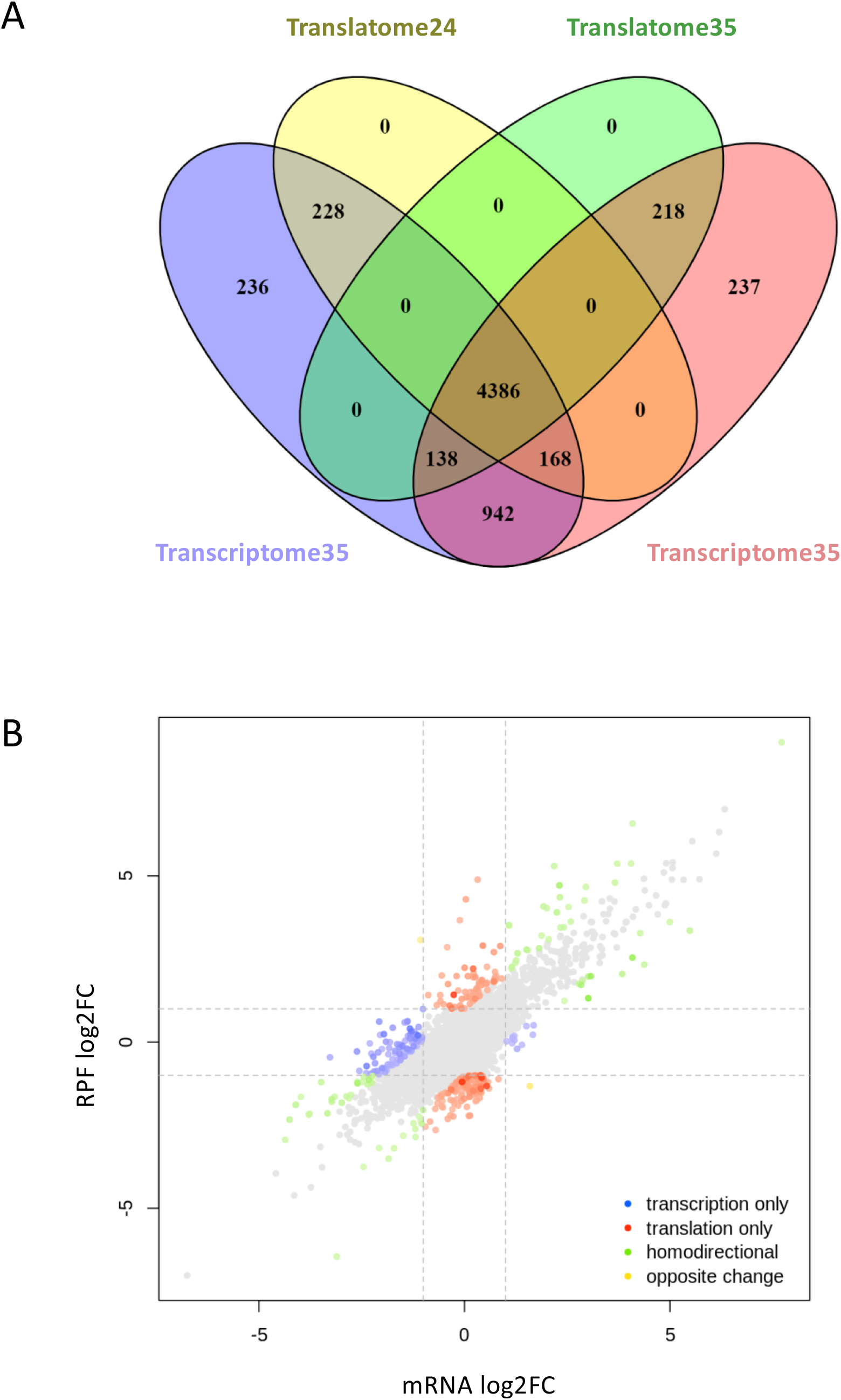
Comparative analysis of germinated pollen transcriptomes and translatomes at 24°C and 35°C. A, Venn diagrams were drawn with the gene lists of transcriptomes and translatomes at both optimum and restrictive temperatures. B, the graph shows the differential expression analysis correlation obtained with the Xtail pipeline for the transcriptome (RNA log2FC) and the translatome (RPF log2FC) of germinated pollen comparing the treated (35°C) versus the non-treated samples (24°C). The points represent individual genes colour-coded according to their differential values. The intensity of the colour is proportional to the p-value, with darker colours indicating more significant p-value.

To perform a thorough statistically-based analysis of differential expression for the transcriptomes and translatomes at 24°C and 35°C, we used the Xtail pipeline, a DESeq2-based protocol with improved sensitivity and accuracy to quantify differential translations with ribosome profiling data (46). To apply the Xtail analysis, we first generated a valid list of genes using only those present in either the translatome 24 or the translatome 35 to avoid invalid data processing for those genes present in transcriptomes but absent in the translatome gene lists. The valid list of genes for Xtail analysis contains 5138 genes (Supplementary Table S3). Then we run the Xtail pipeline with the valid list of genes to obtain a table (Supplementary Table S4) that summarizes the quantitative differences at the transcriptional (log2FC mRNA) and translational level (log2FC RPF). Figure 5B displays the plot of transcriptional and translational differences of *A.thaliana* pollen germinated *in vitro* upon heat shock. We found that the vast majority of genes with differential expression (log2FC > 1 or log2FC < −1) display homodirectional changes for the transcriptome and the translatome, with a low number of genes displaying unique transcriptional or translational changes (blue and red coloured dots in Figure 5B). We initially picked those up-or down-regulated genes, both at the translational and transcriptional level, as a proxy for the major increases and reductions of protein levels. According to the restrictive selection criteria we found 355 up-regulated and 300 down-regulated genes (Supplementary Table S4). Both lists of genes were analysed with the g:Profiler tool for the presence of enriched terms compared in this case to the valid list of 5138 genes used for the Xtail analysis.

#### Heat up-regulated HSP and ER stress genes

As it is shown in Figure 6A among the up-regulated genes we could find enriched terms related to ‘response to heat’, ‘protein folding’, ‘response to temperature stimulus’, ‘unfolded protein binding’ and ‘endoplasmic reticulum’. Figure 6B shows the comparison of RNA and RPF log2FC values > 1 for a list of representative genes of the most significant GO terms identified, including members of the large heat-shock protein familiy mostly of the small HSP20-like, DNAJ chaperones, ER-related proteins, and other members of the unfolded-protein-response (UPR) and the cytosolic-protein-response (CPR) pathways (Supplementary Table S5). In a more detailed analysis, Table 2 shows the transcriptional and translational expression levels in germinated pollen for the list of 13 genes co-induced by the three stressor pathways (heat shock, UPR and CPR), previously reported for *A.thaliana* leaves (69). It is remarkable that all the genes are also expressed and induced in germinated pollen upon heat-stress, with the only exception of *AtbZIP60*, a key gene regulator of the UPR pathway (70), present in germinated pollen although neither transcriptionally nor translationally induced by heat. Altogether the data show that the highly conserved and protective cellular response to heat stress in terms of protein folding seems to be activated in pollen as in vegetative tissues, sharing transcriptional signatures with the related UPR and CPR pathways.

**Table 2.**
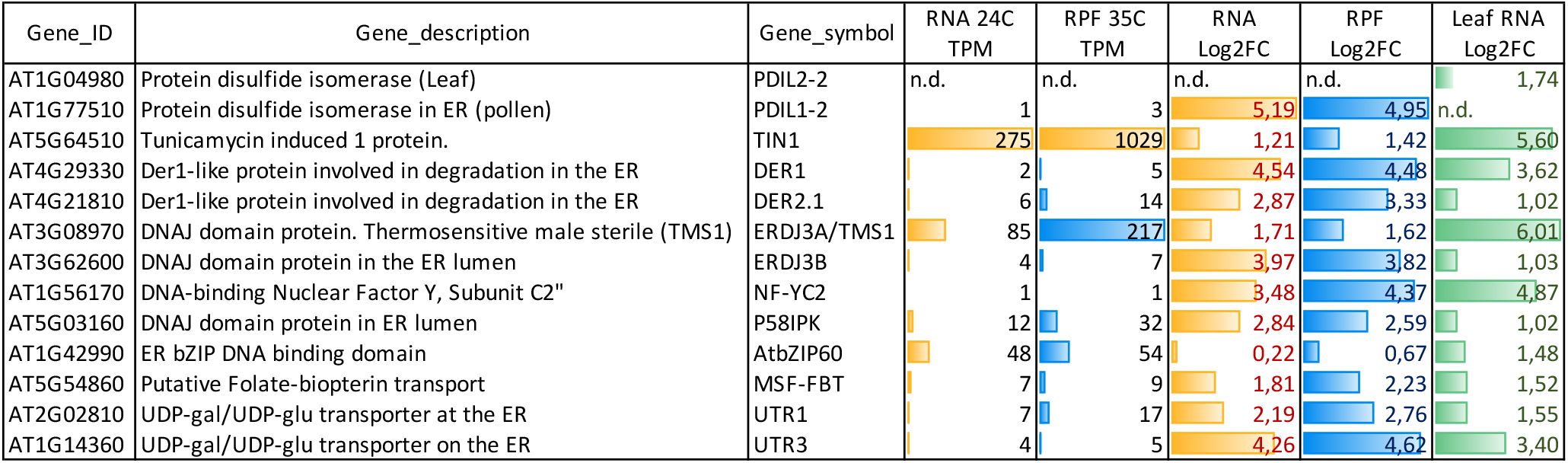
Key genes involved in heat stress and UPR in leaf are UP-regulated in pollen. Key genes involved in heat stress, UPR and CPR pathways in leaf and pollen. Data include gene expression data for 13 genes co-induced in plant leaves by different stressors as previously published, and the corresponding data from this work. We include pollen RNA and RPF values at 24°C, and their log2FC (35/24) from our dataset.

**Figure 6.**
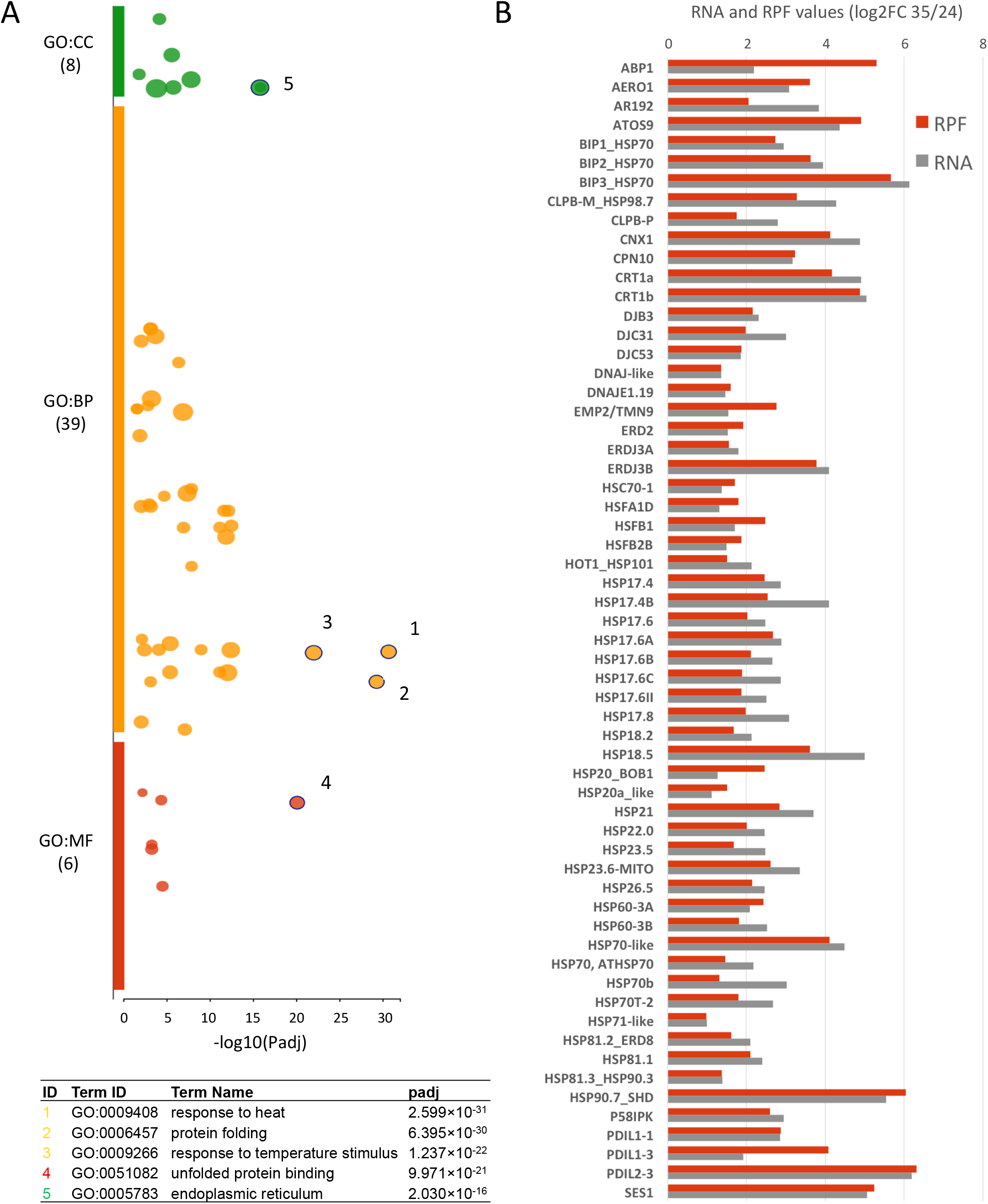
GO enrichment of up-regulated genes at high temperature. A, the g:Profiler tool was used to uncover GO term enrichment of induced genes both at the transcriptional and the translational level, identified through the Xtail analysis. The dot size is proportional to the number of genes included in the term, and the most significant terms are highlighted. The GO enrichment analysis was performed against the custom list of 5138 valid genes of the Xtail analysis. B, shows the graphical representation of log2FC RNA and RPF values for a selected list of genes from the most significant GO terms sorted by gene families.

#### Heat down-regulated mainly transport genes

On the other hand, we examined the down-regulated list of genes with the aim to identify heat-sensitive pathways. We found strong enrichment of terms defined as ‘secondary active transmembrane transporter’, ‘active ion transmembrane transporter’, or ‘carbohydrate:proton symporter’ all linked to key pathways related to ion homeostasis and secondary transport of carbohydrates coupled to H^+^ gradient (Figure 7A). Figure 7B shows the comparison of RNA and RPF log2FC values < 1 for transporter genes detailed in Supplementary Table S6. One paradigmatic case is the enrichment of the cation/H^+^ exchangers (CHX) down-regulated upon heat stress. The *CHX* gene family contains 28 members in *A.thaliana* which have been postulated to participate in diverse pollen developmental activities such as ion and metabolite transport, osmotic adjustments, vacuole formation and vesicular trafficking among others (71). Although some functional redundancy has been found between CHX transporters (72–74), according to our data shown in Table 3, 23 of them displayed > 2 TPM values in both 24°C and 35°C transcriptomes and, therefore, can be considered as *bona fide* members of the germinated pollen transcriptome at permissive and restrictive temperatures. Data presented in Table 3 show a global negative effect of temperature on CHX levels, since most of the gene family members display a down-regulation both at the transcriptome and translatome levels although only 7 members display values below our stringent cutoff (log2FC < 1) as shown in Figure 7B. In addition to the *CHX*s, another gene family negatively affected by elevated temperature at the transcriptional and translational level during pollen germination is the sucrose/H^+^ transporter family. Arabidopsis has 9 genes encoding sucrose/H^+^ symporters (*AtSUC1-AtSUC9*) mostly expressed in sink tissues and phylogenetically distributed in three subtype groups (75). Our transcriptome and translatome data (shown in Table 3) indicate that 5 out of the 9 gene family members are transcribed and 4 of them properly translated at optimum temperature, but most of the members display negative log2FC values for transcriptome and translatome. In fact, two of them (*AtSUC7* and *AtSUC8*) show the top-ranking values of down-regulated genes by heat stress according to the results of the Xtail analysis (Supplementary Table S4). In a similar manner, the related family of hexose transporters STP is negatively affected by temperature with 3 members displaying negative log2FC values < 1. In addition to these predominant gene families, the list of membrane transporters down-regulated at high temperature is remarkable including primary membrane-energizing transporters such as AHA 6 which may be highly deleterious for pollen tube growth considering the importance of H^+^ homeostasis. In contrast, the vacuolar H^+^ ATPase VHA, and many diverse membrane transporters showed little to no significant changes or even induction after the heat shock (Figure 7B). Overall, these data suggest that a failure in the orchestrated presence of at least 39 membrane transporters under heat stress could be detrimental for proper germination and growth of the pollen tube.

**Table 3.** Gene epxression levels for *CHX* and *SUC* genes expressed in germinated pollen. Riboprofiling data for gene families of transporters with altered expression. Transcriptome and translatome data (mean TPM values), together with differential expression analysed with Xtail pipeline is shown for different gene members of the cation/ H^+^ (CHX) and sucrose/ H^+^ (SUC) transporters. NA means non-available data in the Xtail analysis.

| Gene_symbol | Gene_ID | 24°C |  | 35°C |  | XTAIL data |  |
| --- | --- | --- | --- | --- | --- | --- | --- |
|  |  | RNA (mean TPM) | RPF (mean TPM) | RNA (mean TPM) | RPF (mean TPM) | RNA log2FC | RPF log2FC |
| ATCHX23 | AT1G05580 | 4818,7 | 2518,1 | 4001,0 | 1972,7 | -0,1 | -0,4 |
| ATCHX8 | AT2G28180 | 510,0 | 993,6 | 318,0 | 866,0 | -0,6 | -0,3 |
| ATCHX5 | AT1G08150 | 466,2 | 537,1 | 243,5 | 275,1 | -0,8 | -1,0 |
| ATCHX6A | AT1G08140 | 387,4 | 373,8 | 259,0 | 253,8 | -0,5 | -0,6 |
| ATCHX19 | AT3G17630 | 196,8 | 218,9 | 65,5 | 102,4 | -1,5 | -1,2 |
| ATCHX9 | AT5G22910 | 119,8 | 246,8 | 68,1 | 252,3 | -0,7 | 0,0 |
| ATCHX2 | AT1G79400 | 98,9 | 77,8 | 99,8 | 98,8 | 0,1 | 0,3 |
| ATCHX26 | AT5G01680 | 95,4 | 138,9 | 27,6 | 74,5 | -1,7 | -1,0 |
| ATCHX15 | AT2G13620 | 88,6 | 197,6 | 89,7 | 179,0 | 0,1 | -0,2 |
| ATCHX10 | AT3G44930 | 84,7 | 107,3 | 59,8 | 89,7 | -0,4 | -0,3 |
| ATCHX11 | AT3G44920 | 66,2 | 102,5 | 34,1 | 71,7 | -0,9 | -0,6 |
| ATCHX12 | AT3G44910 | 61,5 | 58,3 | 19,7 | 56,7 | -1,5 | -0,1 |
| ATCHX21 | AT2G31910 | 58,2 | 133,2 | 13,3 | 53,1 | -2,0 | -1,4 |
| ATCHX3 | AT5G22900 | 54,9 | 143,4 | 16,7 | 64,1 | -1,6 | -1,2 |
| ATCHX4 | AT3G44900 | 53,2 | 78,7 | 14,7 | 33,5 | -1,8 | -1,3 |
| ATCHX6B | AT1G08135 | 40,9 | 83,1 | 21,0 | 67,9 | -0,8 | -0,4 |
| ATCHX7 | AT2G28170 | 28,9 | 28,2 | 5,1 | 18,0 | -2,4 | -0,7 |
| ATCHX28 | AT3G52080 | 21,8 | 52,0 | 31,2 | 92,5 | 0,6 | 0,8 |
| ATCHX27 | AT5G01690 | 12,9 | 4,4 | 4,7 | 1,9 | -1,3 | -1,3 |
| ATCHX14 | AT1G06970 | 10,5 | 26,3 | 4,3 | 9,4 | -1,1 | -1,6 |
| ATCHX13 | AT2G30240 | 3,9 | 4,0 | 5,2 | 4,3 | 0,5 | 0,0 |
| ATCHX25 | AT5G58460 | 2,7 | 0,7 | 3,2 | 0,5 | NA | NA |
| ATCHX18 | AT5G41610 | 2,2 | 0,6 | 2,2 | 1,0 | NA | NA |
| ATSUC9 | AT5G06170 | 258,5 | 621,4 | 99,1 | 432,4 | -1,3 | -0,6 |
| ATSUC7 | AT1G66570 | 11,6 | 14,0 | 0,1 | 0,1 | -6,7 | -7,0 |
| ATSUC8 | AT2G14670 | 11,2 | 22,5 | 0,4 | 1,7 | -4,6 | -4,0 |
| ATSUC3 | AT2G02860 | 6,9 | 6,1 | 9,0 | 8,8 | 0,5 | 0,4 |
| ATSUC1 | AT1G71880 | 3,5 | 0,5 | 2,9 | 0,4 | NA | NA |

**Figure 7.**
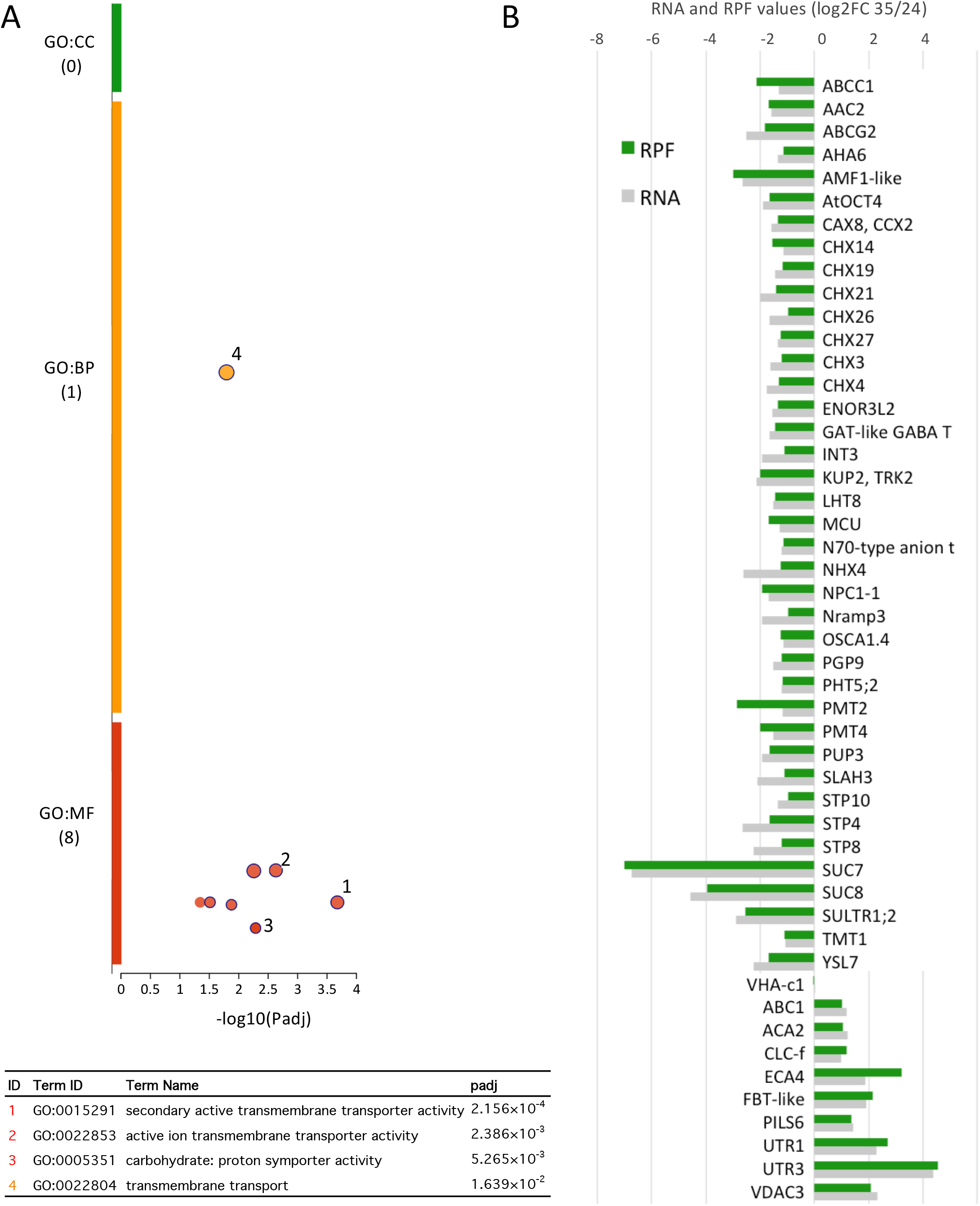
Heat down-regulates mostly membrane transport genes. A, the g:Profiler tool was used to uncover GO term enrichment of down-regulated genes both at the transcriptional and the translational level, identified through the Xtail analysis. The dot size is proportional to the number of genes included in the term, and the most significant terms are highlighted. The GO enrichment analysis was performed against the custom list of 5138 valid genes of the Xtail analysis. B, shows the graphical representation of log2FC RNA and RPF values for a selected list of genes from the most significant GO terms sorted by gene families.

To gain further insight into the roles of the heat-sensitive membrane transporters, we checked their expression during pollen development, as previous studies showed specific expression patterns during pollen maturation (76). We used transcriptome data published by others (57,59) showing expression of *A.thaliana* pollen in different stages: dry pollen, pollen germinated *in vitro* at two different times (0.5h and 4h), pollen germinated *semi in vivo* (*SIV*), and sperm cells. Germinated pollen includes a vegetative tube cell and two sperm cells, and when germinated in *SIV* condition refers to pollen tubes emerged from a cut style on nutrient medium after germinated on a stigma (59). As shown in Figure 8A, we found a striking relationship between a subset of heat-down regulated transporters in our analysis and their up-regulation in *SIV* germinated tubes at basal temperature compared to dry pollen. To further illustrate this finding, we used the web-based tool ‘Arabidopsis Heat Tree Viewer’ to compare 5 different pollen developmental stages. As it can be seen in Figure 8B, there is a notable up-regulation of gene expression for at least 23 transporters after 4 hours *in vitro* growth and especially in tubes grown under *SIV*. We also checked the expression during these 5 pollen stages of the heat-induced genes identified previously in our dataset, to find co-regulated patterns, and we focused on the 17 pollen-expressed genes induced by heat belonging to the conserved family of the small heat shock proteins (sHSPs) in plants (77,78). We found that 13 of them displayed a similar pollen gene expression pattern to the transporters, with increased expression during *in vitro* germination further enhanced in *SIV* conditions, whereas in shoots, although highly induced by heat, their basal expression is null (Figure 8C). These data suggest that during pollen germination there is a strong requirement for expression of key ion and carbohydrate transporters together with proteins involved in folding and refolding, and, during heat stress conditions, in spite of a proper induction of the folding machinery, a lack of gene expression response for the transporters may lead to pollen tube growth defects.

**Figure 8.**
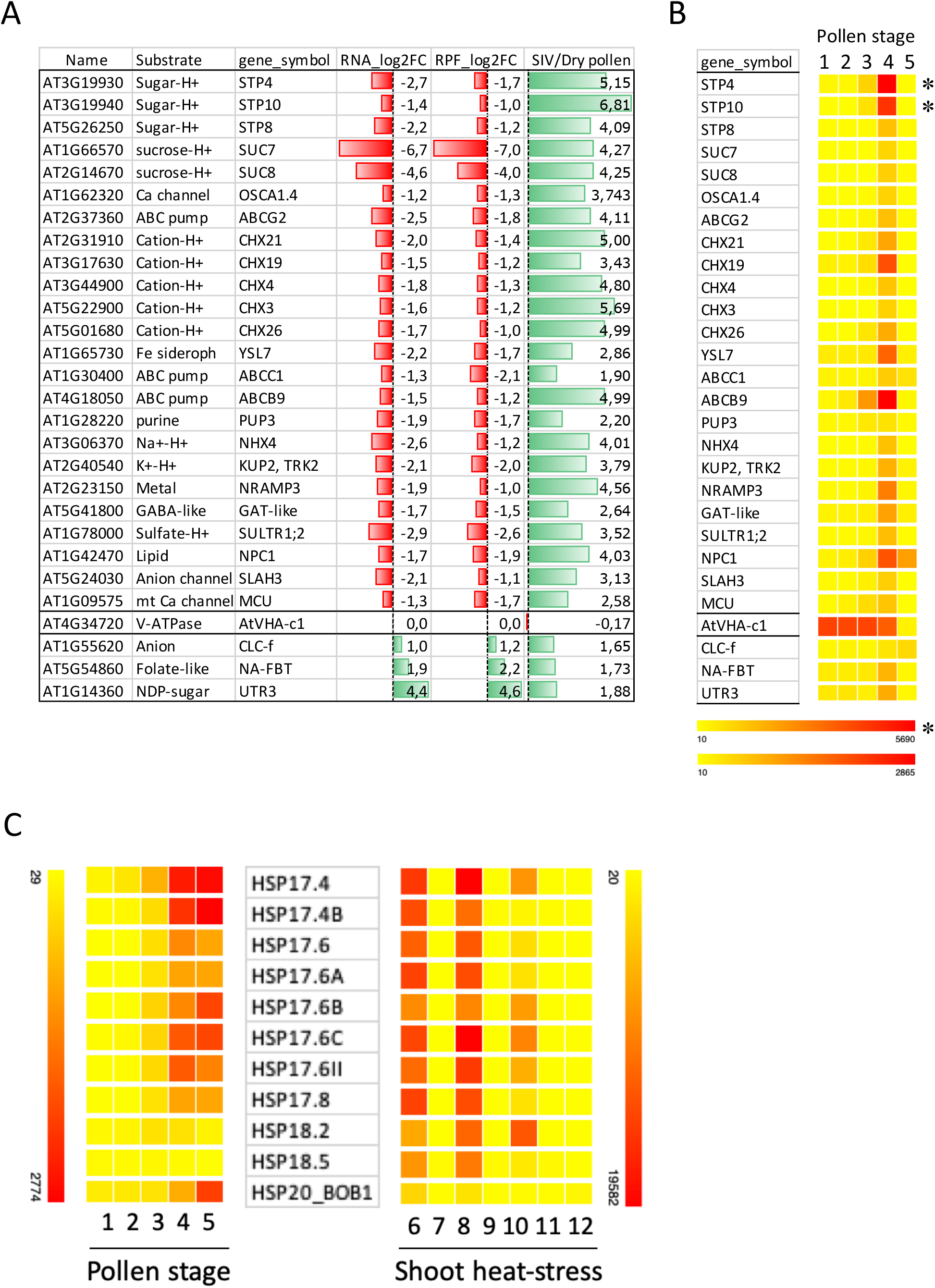
Down-regulated transporter genes display opposite trend during *SIV* growth. A, the list of down-regulated membrane transporter genes was scored for their gene expression value ratios of SIV/dry pollen showing reverse tendency to heat-response. B, the list of down-regulated membrane transporter genes was analysed with the ‘Arabidopsis Heat Tree Viewer’ web tool to represent gene expression according to values indicated in the scales, either with or without asterisk. C, 17 sHSPs expressed in pollen were analysed similarly with the ‘Arabidopsis Heat Tree Viewer’ web tool and the scales showed below. Numbers indicate different pollen developmental stages: 1, dry pollen; 2, 0.5h *in vitro* pollen tubes; 3, 4h *in vitro* pollen tubes; 4, *SIV* pollen tubes; 5, sperm; 6, heat 1h shoot; 7, heat_control 1h shoot; 8, heat 3h shoot; 9, heat_control 3h shoot; 10, heat 4h shoot; 11, heat_control 4h shoot.

#### Changes in translational efficiency during heat stress

We analysed whether Riboprofiling revealed any alteration in translational efficiency (TE), referred to the changes in the ratio of RPF to RNA counts for a given gene between two different conditions. The Xtail algorithm uses two parallel pipelines to quantify the TE changes, first as the difference between the log2FC of RPF and RNA across the two temperature conditions, and second as the difference between the log2 ratios of RPF to RNA in both conditions (log2Rs). Both calculations should yield a similar result, and the more conserved pipeline with better p-value is selected for the final assessment (log2FC_TE) of differential translation (46). Figure 9A shows the correlation plot of log2Rs at 24°C and 35°C where a notable homodirectional pattern, similar to the transcriptome versus translatome shown in Figure 5, can be observed. In spite of some exceptions (coloured dots in Figure 9A), most of the alterations in TE occur for RNA and RPF in a correlated manner, therefore we see no evidence of wide effects of high temperature on translation dynamics. According to acceptable p-value < 0.05 we found 77 and 120 genes with down-regulated or up-regulated log2FC_TE values respectively (Supplementary Table S4). In spite of the low number of genes affected (3.8%), we investigated the nature of those changes by considering all possible regulatory scenarios. Figure 9B shows a summary of all the 9 possible changes at either the RNA or RPF between both temperatures that may lead to final TE changes. As it can be seen, among the down-regulated TE genes, a large number (58.4%) show changes caused by RPF with no RNA alterations, suggesting specific translational inhibition mechanisms. Intriguingly, the case is the opposite for a large proportion (43.3%) of the up-regulated TE genes with most of the changes affecting at the RNA level. The graphical representation of log2FC_TE as a function of the p-value is shown in the volcano plot of Figure 9C. Some exceptions to the homodirectional pattern (highlighted in Figure 9C) are the ATP-dependent caseinolytic (Clp) protease (AT1G09130) which suffers a strong down-regulation at the translational level with the increase in temperature, or the annotated as non-coding RNAs AT5G06845 and AT1G05853 with surprisingly enhanced ribosome footprints at 35°C. In fact, the top three genes with up regulated log2FC_TE values are non-coding RNAs with RPF values unexpectedly increased at 35°C (Supplementary Table S4). The functional interpretation of RPF reads on annotated non-coding RNAs remains to be clarified and is considered in the next section.

**Figure 9.**
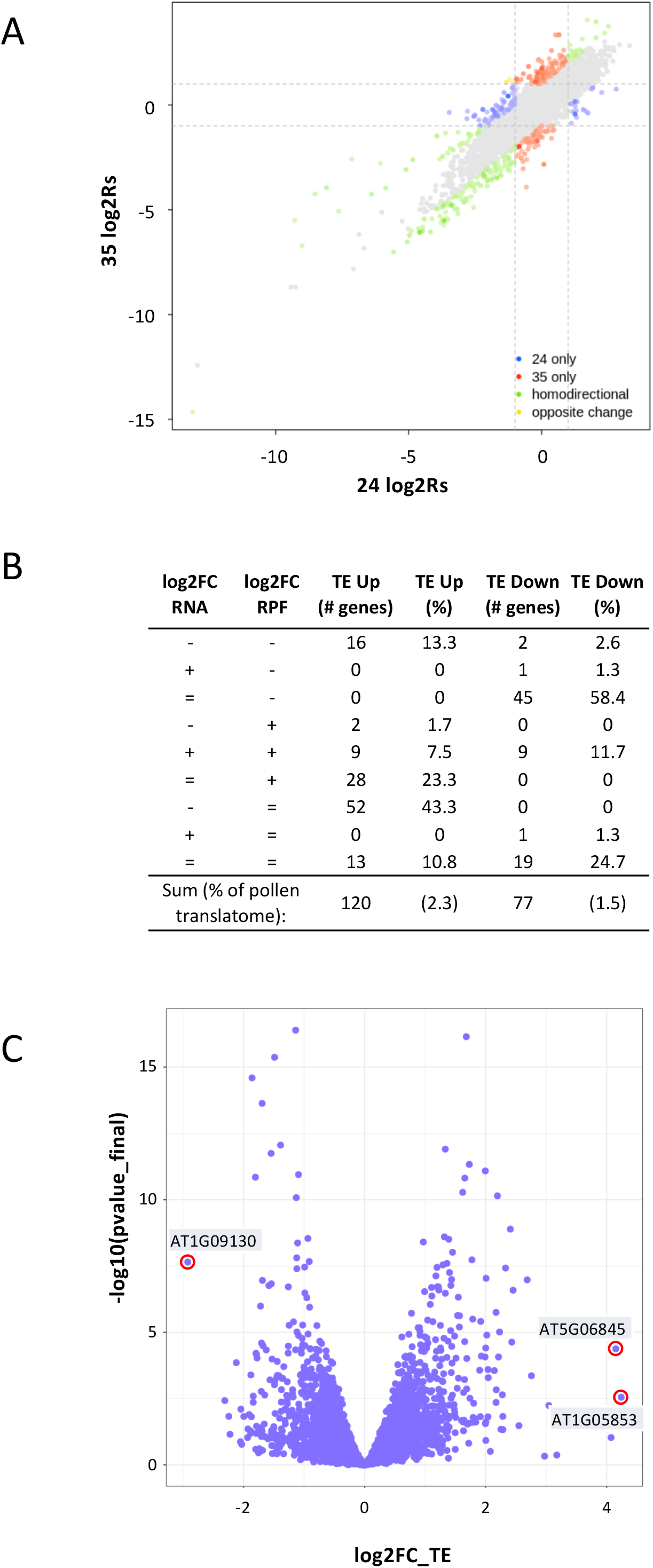
Analysis of changes in translational efficiency (TE). A, the plot shows the correlations of the translational efficiency data (log2Rs) of germinated pollen at 24°C and 35°C obtained with the Xtail pipeline. B, the table shows the different scenarios that account for TE changes. Symbols (−), (+) and (=) indicate a reduction, increase or no change in gene expression, respectively. The numbers are relative to the total number of differentially expressed genes (5138). C, volcano plot of differential translational efficiency (log2FC_TE) comparing high (35°C) and optimum temperature (24°C) as a function of the p-value significance. Some examples of significantly enhanced or reduced translational efficiency are highlighted with red circles.

### Revisiting predicted functions and annotations for the translatome of germinated pollen

In spite of experimental and bioinformatics removal of rRNAs, tRNAs, snRNAs and snoRNAs, our transcriptome and translatome lists contained genes annotated as non-coding RNAs. These include mostly long-non-coding RNAs, transposable elements and a few small-nucleolar RNAs that probably escaped the filtering due to annotation issues. Apart from their possible roles as regulatory transcripts with transcriptional and post-transcriptional impact, some of these RNAs show significant RPF reads suggesting the presence of ribosomes according to the very strict RPF size used in the analysis (only 27 and 28 nt long). The advantage of performing Riboprofiling studies is that the parallel analysis of the transcriptome versus the translatome allows the detailed examination of previous annotations that may need revision. On the one hand, the use of the Integrative Genomics Viewer (IGV), with graphical visualization of the read coverage, allows the fine observation of the mapped genes (50). On the other hand, to make sure that RPFs derive from RNase protection by the ribosomes, we can exploit the 3-nt periodicity of the ribosome to confirm the presence of ribosomes in non-coding RNAs. To this extent we have used the Ribowave pipeline to examine the periodicity of PF P-sites of several examples of non-coding RNAs presenting detectable RFPs.

One paradigmatic example is the gene AT2G41310 with very high transcriptome values both at 24°C and 35°C, but very low RPF values (Supplementary Tables S3 and S4). This gene, annotated as ARR8/ATRR3, encodes an A-type response regulator involved in cytokinin-mediated signaling, and it caught our interest as it was not expected to be expressed in pollen. Figure 10A shows the IGV coverage data showing that the reads fall into the 3’ untranslated region overlapping with a non-coding RNA (AT2G09250) in the opposite orientation. Therefore, there is no doubt that read coverage in that region does not correspond to the encoded ARR8 protein. We then used the Ribowave pipeline to examine the periodicity of PF P-sites in that region. The results shown in Figure 10B demonstrate the presence of periodic P-site footprints at both temperatures concentrated in one of the three reading frames. Although the density of PF P-sites and the coverage values are rather low compared with normally translated genes (compare with the examples shown in Figure 4B), the data suggest that this gene could be translated in germinated pollen. Similar observations were found for other non-coding-RNAs showing detectable RPF reads in the genes AT5G06845 and AT1G05853, however the limiting PF P-site density and low coverage values in most of the cases precludes the achievement of any definitive conclusion as to whether those genes are indeed translated.

**Figure 10.**
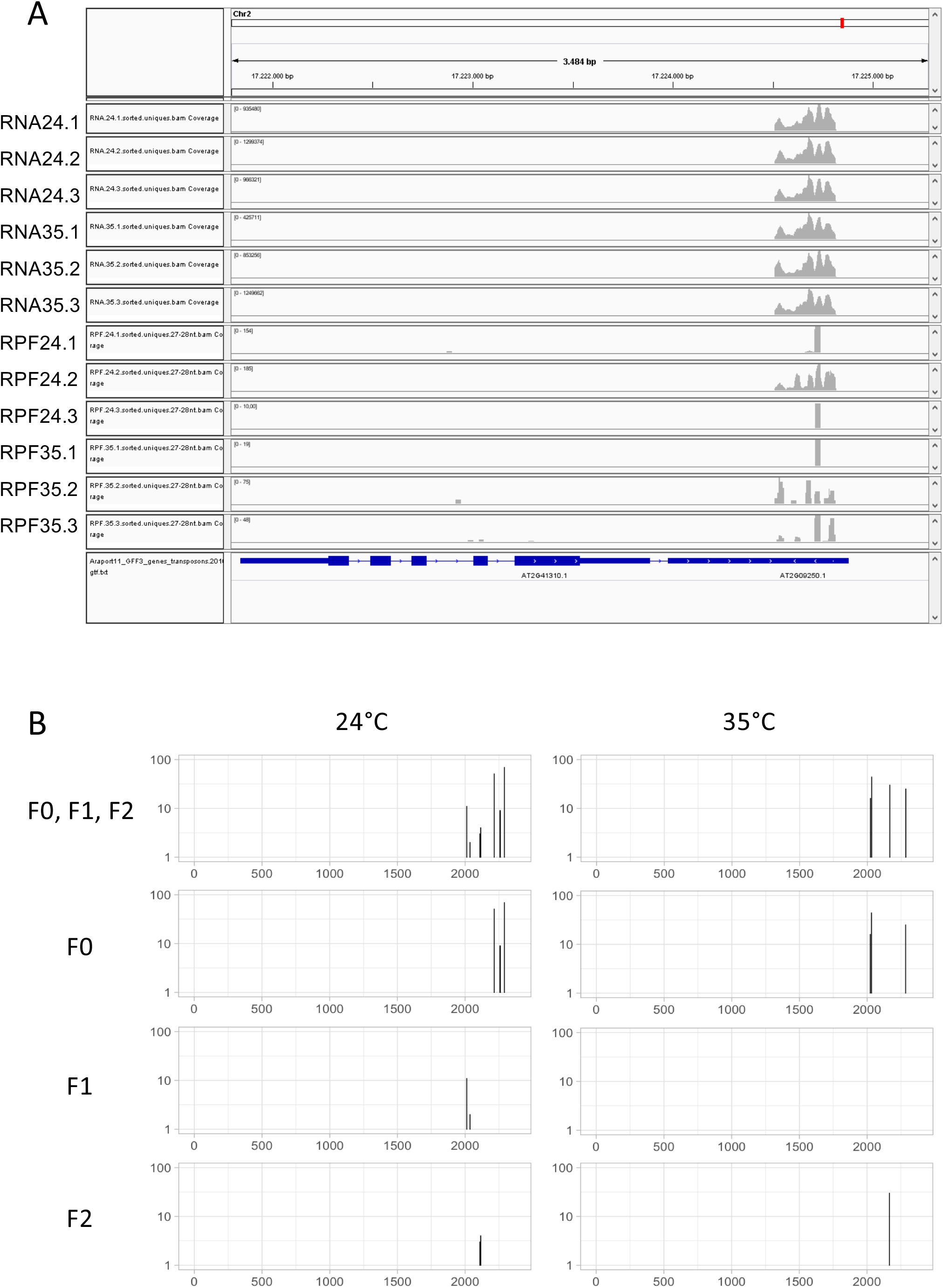
A case of problematic gene annotation and singular translational profile. A, the IGV visualization of mapped reads on the gene AT2G41310 shows an overlapping of the reads at the 3’ UTR where a non-coding RNA gene (AT2G09250) is located in the antisense orientation. B, the periodicity of the PF P-site distribution for the mapped reads shown in Panel A was computed in the 3 reading frames (F0, F1, F2), or for every reading frame separately, showing periodic signal only in the region coincident with AT2G09250.

In this work we have focused on annotated sequences, but we have further exploited the results of our Riboprofiling experiments to help characterize the translational status of improperly annotated pollen genes, by using a different alignment pipeline of our RPF libraries that avoids the removal of non-coding RNAs, as shown in another study in collaboration with G.Miller (submitted manuscript). Altogether, the data of both manuscripts highlight the idea that additional efforts are needed to help improve the genome annotations of a well-curated reference genome as it is the case of *A.thaliana*.

## DISCUSSION

In this work we have used, for the first time, a high-resolution ribosome profiling technology to achieve a comprehensive study of how heat affects both the transcriptome and the translatome of *A.thaliana in vitro* germinated pollen. From the different cell types present in mature pollen (2 sperm cells and one vegetative cell), it is the vegetative cell the one involved in pollen tube growth and the most active cell during pollen germination, as its mission will be the safe delivery of the sperm cells into the ovule to accomplish the double fertilization process. Thus, the transcriptome and translatome changes shown in this work are largely affecting the vegetative cell, and in fact we have observed strong translational repression of GO terms specific for sperm cells. Our results provide novel insights on the effects of high temperature on pollen gene expression after obtaining a complete dataset of the transcriptome and the translatome of *in vitro* germinated pollen at basal (24°C) and restrictive temperature (35°C). The differential expression analysis uncovered new findings that can be summarized as follows: (i) the conserved cellular machinery to cope with heat stress is well functioning in germinated pollen, as we found up-regulated many genes of the heat-response, the UPR and the CPR pathways; (ii) the enrichment of membrane transporters, involved in ion and carbohydrate transport among the down-regulated genes, identified heat-sensitive targets; (iii) we detected a co-regulation between heat-sensitive membrane transporters and heat-induced proteins during specific stages of pollen germination, namely *in vitro* and *SIV*, illustrating new co-regulatory aspects of pollen germination; (iv) although we found a high correlation between transcription and translation, specific translational efficiency alterations could also be uncovered in germinated pollen; and (v) we could detect the presence of ribosome footprints on non-coding RNAs, illustrating the need of further refinement for gene expression on germinated pollen.

All these relevant data could be obtained by means of the Riboprofiling technology, a powerful technological advance that allows the collection of genome-wide information of ribosome footprints with sub-codon resolution in a quantitative manner. One advantage of the Riboprofiling is that simultaneous analysis of RNAseq fragments allows the computation of translational efficiency for the whole transcriptome set. The results of the three biological replicates for basal (24°C) and restrictive (35°C) libraries verified the reproducibility of the experiments, and the use of DESeq2-based statistical tools, to characterize the differential analysis, provided robust results in terms of statistical significance. We used a very stringent threshold value (2 TPM) as a conservative gene expression cutoff for the benefit of achieving a reliable dataset, which seemed appropriate as compared to previously published pollen transcriptomes.

An important question is about the nature of the up- and down-regulation gene expression patterns. As we have monitored RNA and RPF levels in response to heat, it is unclear whether their alterations are due to changes in the synthesis or degradation, or both. Pollen grains contain stored mRNA, and it is assumed the mRNAs can be released and immediately translated after pollination to support rapid tube growth (79). Recent data have shown the presence of cytoplasmic mRNA granules, namely stress granules (SGs), in mature pollen grains co-localizing with processing body (PB) proteins, thus suggesting the presence of PBs during pollen maturation (80). Both SGs and PBs are intimately connected with each other and with the translational machinery, to modulate the mRNA turnover during stress conditions (81). Thus, an increase in transcript could reflect a combination of newly-synthesized transcripts and released mRNAs from storage granules, and a down-regulation could be caused by reduced transcription or enhanced degradation in PBs, or both.

The differential gene expression analysis provided clues on the cellular pathways up-regulated and down-regulated by heat stress. Among the up-regulated pathways, we could find a significant enrichment of GO terms related to heat shock response, protein folding and quality control ER-dependent pathways (Figure 6 and Supplementary Table 5). In many cases these genes share transcriptional up-regulation with a variety of cellular stresses in plant leaves as shown in Table 2. For instance, we found many up-regulated genes encoding ER proteins as the HSP70 BiP proteins (BIP1, BIP2, BIP3), the DNAJ chaperones (ERDJ3A, ERDJ3B, P58IPK); calnexin and calreticulin proteins (CNX1, CRT1a, CRT1b), disulphide isomerases (PDIL1-1, PDIL1-3, PDIL2-3), and proteins involved in the ER-associated degradation pathway (ERAD) as DER1 and DER2.1. Furthermore, key transcription factors involved in the initiation of these pathways, as bZiP28, bZip60 and NF-YC2 are also expressed in pollen tubes (Tables 2 and S4). These results demonstrate that *A.thaliana* pollen tube is able to sense and respond to heat stress, by specifically up-regulating genes needed for thermotolerance in a similar manner to ER-dependent stress responses in leaves (82). In addition to the ER-dependent folding and quality control machinery, we also found heat up-regulation of a few membrane transporters (Figure 7). For instance, UTR1 and UTR3 are nucleotide sugar transporters required to incorporate UDP-sugars into the ER for glycosylation reactions required in the UPR pathway (83). A putative folate-biopterin transporter that belongs to the major facilitator superfamily (AT5G54860) is also induced by heat, and could be important in tolerance to abiotic stress (84). Additional ion transporters with functions in the ER or Golgi were also induced, as voltage-gated chloride channel (CLC-f) and calcium pumps (ECA4 and ACA2).

As a step to identify the molecular basis of heat sensitivity, we studied enriched GO terms of down-regulated genes (Figure 7 and Supplementary Table 6). We focused on membrane transporters, as they were enriched among the 300 genes down-regulated by heat. Over 260 transporter genes expressed in germinated pollen we found 39 of them (14.8%) down-regulated, so for the majority of transporters, including the vacuolar H^+^ pump ATPase VHA, heat had little effect. However, down-regulation of those 39 genes affected to transporters with key functions during pollen tube growth, including the primary pump H^+^-ATPase AHA6, calcium channels like OsCA1.4, ABC pumps and H^+^-coupled secondary transporters for K^+^, anions (sulfate, phosphate), metals, and amino acid transporters like LHT8. Most prominent is the large number of H^+^-coupled sugar transporters (*SUC* and *STP* genes) and members of the cation/H^+^ exchanger family (*CHXs*) with most of their family members affected. The multiplicity of *CHX* genes in pollen function is only partially understood. Based on previous studies CHX16-20 behave like a cation/ H^+^ exchanger that is proposed to mediate pH changes locally and transiently across endomembranes and/or plasma membrane (85). Triple mutants *cxh 17/18/19* showed a disorganized architecture of the pollen wall, reduced male fertility and seed set (74). CHX21/23 transporters are important for pollen tube targeting to the ovule as the *chx21/23* double mutant pollen fail to reach the ovule and are male sterile (72). Although the specific functions of additional *CHX* genes in pollen remain to be characterized, it is likely they participate in critical functions for tube growth, including signal transduction, tube navigation, and targeted delivery of materials to the wall or the exterior, events that ensure male reproductive success. Altogether, the massive down-regulation of membrane transporters by heat may lead to devastating consequences due to the strong requirements of highly active vesicular transport during pollen tube growth. We hypothesize that such fatal depletion of key transporters closely linked to pH regulation, signalling, nutrients and osmotic adjustment may be the main cause of defective pollen growth under elevated temperature conditions.

We also discovered that about two thirds of these down-regulated transporters by high temperature are also up-regulated during *in vitro* germination and especially when pollen tubes were grown under *semi in vivo* (*SIV*) conditions (59). For instance, *SIV* conditions enhanced expression of *SUC7/8* and *STP4/8/10* by 16 o 64-fold relative to dry pollen. Similarly, expression of several CHX transporters (*CHX3/4/19/21/26*) are increased 16 to 32-fold under *SIV* conditions. These results suggest that a subset of transporters is sensitive to chemical and physical cues received by pollen from the stigma, or the transmitting tissue or both. Increased expression of sucrose and hexose transporters likely provides carbon source for energy and for synthesizing pollen walls, but also signalling roles have been proposed (86). So, is there any relationship between down-regulation of transporters by heat and their up-regulation by germination conditions? We have shown in Figure 6 and Table 2 that, as in leaves, germinated pollen stressed by heat also triggers up-regulation of folding (chaperone HSPs) and quality control pathways (UPR, CPR, ERAD), and many of those up-regulated genes are also induced under *in vitro* and in *SIV* germination conditions (Figure 8). One prevalent example is the family of cytoplasmic small HSPs (sHSPs), abundantly expressed in germinated pollen under heat stress as they are required to prevent protein aggregation. We have found that 13 out of 17 pollen expressed sHSPs are also induced during *SIV,* thus uncovering potential roles of these chaperones in the germination process. One of these chaperones, the sHSP BOB1 protein, was found incorporated into heat shock granules (HSGs) at high temperature (87). Therefore, we cannot discard that assembly of HSGs, stress granules and processing bodies may lead to either sequestration or targeted degradation of specific mRNAs instead of inhibition of transcription, to explain the gene down-regulation of transporters upon heat stress in germinated pollen.

An advantage of the Riboprofiling technology is that simultaneous identification of mRNA and RPFs allow the quantisation of changes in translational efficiency (TE). The differential analysis of TE upon heat stress revealed that only a minor subset of genes displayed significant TE alterations. As it was shown in Figure 5 the homodirectional changes for RNA and RPF is the prevalent feature and it is also evident when computing the TE changes (Figure 9). The sources of TE alterations are very diverse since they may affect to either RNA or RPF levels both at 24°C and 35°C. It is remarkable that, among the TE down-regulated genes, most of them include changes exclusively in the RPF, as an indication of translation-specific repression. On the other side, among the TE up-regulated genes, almost half of them are caused by changes in RNA levels. The nature of the translational regulations remains to be elucidated.

In addition to detecting gene expression changes, Riboprofiling analysis allows the direct observation of the precise mapping positions of the reads, using integrated visualization programs such as IGV. For instance, a questionable annotation could lead to erroneous hypothesis in the case of the ARR8 encoded gene (AT2G41310) shown in Figure 10. Another example is the case of *AtSUC7,* initially considered as a pseudogene (88), and recently shown to be translated in heterologous systems with functional restrictions on sucrose analogue transport (89). Here we show, for the first time, that *AtSUC7* is transcribed and translated in germinated pollen and it suffers a dramatic down regulation at both transcriptional and translational levels upon heat stress (Table 3).

The Riboprofiling of *A.thaliana* germinated pollen has also uncovered the unexpected presence of RPF reads on several non-coding-RNAs, such as long-non-coding-RNAs, small-nucleolar-RNAs, and transposable elements. In spite of the relevance of this information, the quantification of translation is a problematic issue for small RNAs as the accumulation of the reads in a short sequence may cause artefactual quantifications, especially since most of the cases include genes with very low expression values. Therefore, any interpretation of translation on these kinds of genes should be taken with caution. To corroborate the quantitative RPF data, we have used the Ribowave tool as an additional input to estimate the periodicity of P-sites. Although the lack of well-defined translational start and stop sites makes it difficult for the algorithm to define peptide predictions, it is still possible to extract the PF P-site data and plot it to detect periodic signals as a proxy of ongoing translation. We have studied in detail the case of AT2G09250, a small non-coding RNA overlapping with the 3’ untranslated region of AT2G41310 encoding the ARR8 protein. In this case, the analysis of the periodicity of the PF P-sites as a proxy for the 3-nt ribosomal advance, suggests that this small RNA may be translated. Additional experimental and functional data will be required to unequivocally show that those non-coding-RNAs are indeed subjected to translation during pollen germination and growth.

As a conclusion we can affirm that the use of *in vitro* germinated pollen is a useful system to understand the molecular basis of heat-induced responses. Riboprofiling has provided many answers to better understand pollen gene expression responses, as for instance the particular sensitivity of transporters, at the gene expression level, to the heat insult. Our gene expression data, combined with previously reported information on *SIV* germinated pollen, suggests that pollen tube growth is fully armed, with the co-induction of heat-chaperone proteins and transporters, to cope with a high demand of vesicle transport and resilience to environmental and/or female cues during its journey to the ovule. However, it has also raised many questions that remain to be addressed like the nature of the regulatory molecular mechanisms involved and the translation of annotated genes as non-coding RNAs. The implementation of this powerful technology to the diverse pollen developmental stages and environmental cues in future studies, will provide a more detailed picture of this key aspect of plant biology as it is the fertilization process in a constantly changing environment.

## DATA AVAILABILITY

Raw sequences derived from the Riboprofiling analysis and table containing counts and normalized TPM values for all *A. thaliana* genes have been deposited in the Gene Expression Omnibus database (http://www.ncbi.nlm.nih.gov/geo/) with the accession number GSE145795.

## SUPPLEMENTARY DATA

Supplementary Data are available online.

## ACKNOWLEDGEMENTS

The authors thank the Bioinformatics Core Service of the IBMCP for technical assistance. We also thank José M. Alonso, Anna Stepanova, René Toribio and Mar Castellano for sharing protocols with helpful advices on ribosome profiling, and for critical reading of the manuscript. We also thank to Gad Miller and Nick Rutley for sharing data prior to publication. We are indebted to Heven Sze for giving extraordinary input and very useful ideas to improve the quality of the manuscript.

## FUNDING

This research was funded by the Spanish Ministry of Science Innovation and Universities [BIO2015-70483-R, to A.F.].

## CONFLICT OF INTEREST

The authors declare that the research was conducted in the absence of any commercial or financial relationships that could be considered as a potential conflict of interest.

## Notes

### Competing Interest Statement

The authors have declared no competing interest.

