## Supplemental Figures and Legends for "Transcriptome and translatome changes in germinated pollen under heat stress uncover roles of transporter genes involved in pollen tube growth": Supplemental Figures Poidevin2020.pdf

#### SUPPLEMENTARY FIGURES LEGENDS

Supplementary Figure S1. Correlation analysis for Riboprofiling libraries. A, principal component analysis showing appropriate separation of different components for the two variables under study (type of library and temperature). B, correlation matrix analysis showing Pearson coefficient R as a colour gradient indicated in the figure.

Supplementary Figure S2. Periodicity graphs for RPF libraries. The periodicity data of the RPF libraries and size lengths ranging from 26 to 30 nucleotides is shown. Colours indicate the 3 different reading frames in blue, green and red.

Supplementary Figure S3. Characterization of transcription versus translation at 24°C. A, the GO term enrichment analysis was performed with g:Profiler for transcripts absent in the translatome at 24°C. MF refers to 'Molecular Function' in red, BP to 'Biological Process' in orange, and CC to 'Cellular Component' in green. The dot size is proportional to the number of genes included in the term. The GO analysis was performed against the *Arabidopsis thaliana* genome. B, the IGV visualization of mapped reads on the gene AT5G16020 shows high number of reads for RNA samples but almost no signal in the RPF libraries. As a comparison the adjacent gene AT5G16010 displays high number of reads both in RNA and RPF libraries.

### Figure S1

A

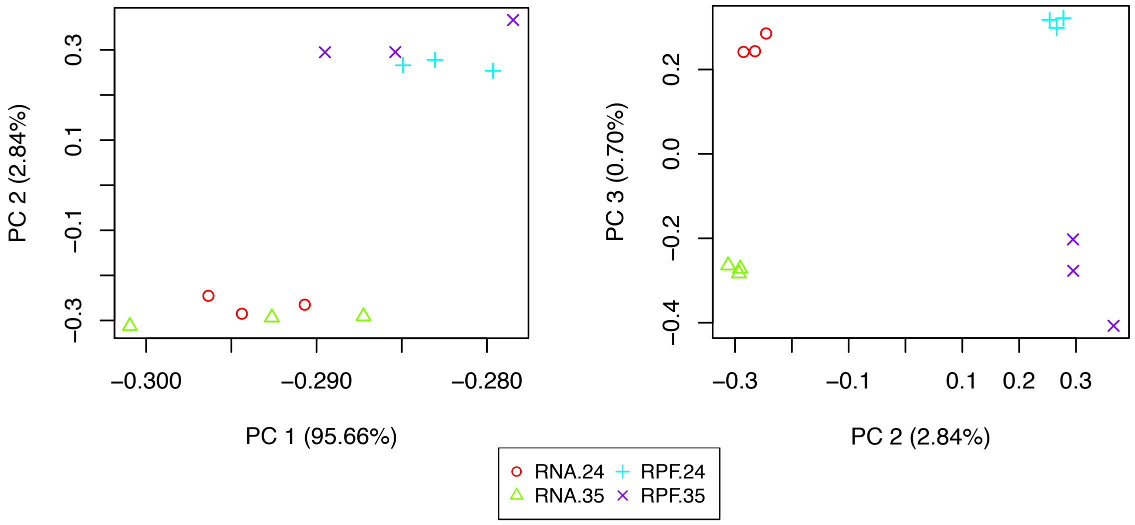
PC 2 (2.84%)
0.3
0.1
−0.1
−0.3
−0.300
−0.290
−0.280
PC 1 (95.66%)
PC 3 (0.70%)
0.2
0.0
−0.2
−0.4
−0.3
−0.1
0.1
0.2
0.3
PC 2 (2.84%)
RNA.24
RPF.24
RNA.35
RPF.35

B

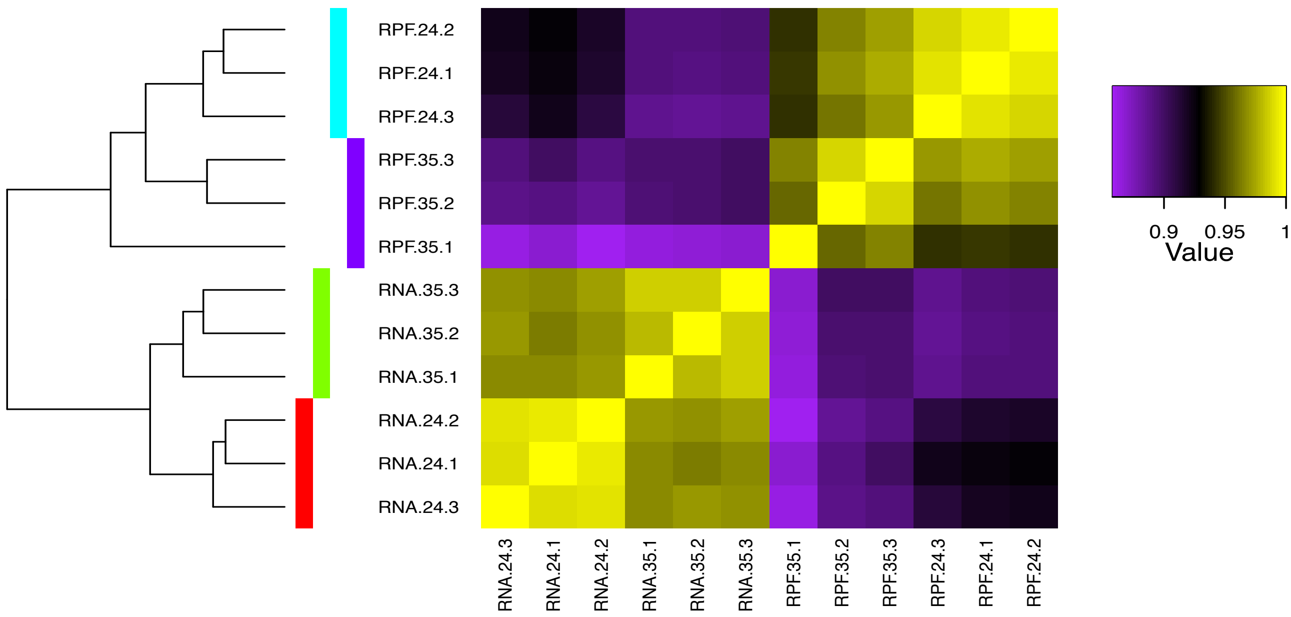
RPF.24.2
RPF.24.1
RPF.24.3
RPF.35.3
RPF.35.2
RPF.35.1
RNA.35.3
RNA.35.2
RNA.35.1
RNA.24.2
RNA.24.1
RNA.24.3
RNA.24.3
RNA.24.1
RNA.24.2
RNA.35.1
RNA.35.2
RNA.35.3
RPF.35.1
RPF.35.2
RPF.35.3
RPF.24.3
RPF.24.1
RPF.24.2
0.9
0.95
1
Value

### Figure S2

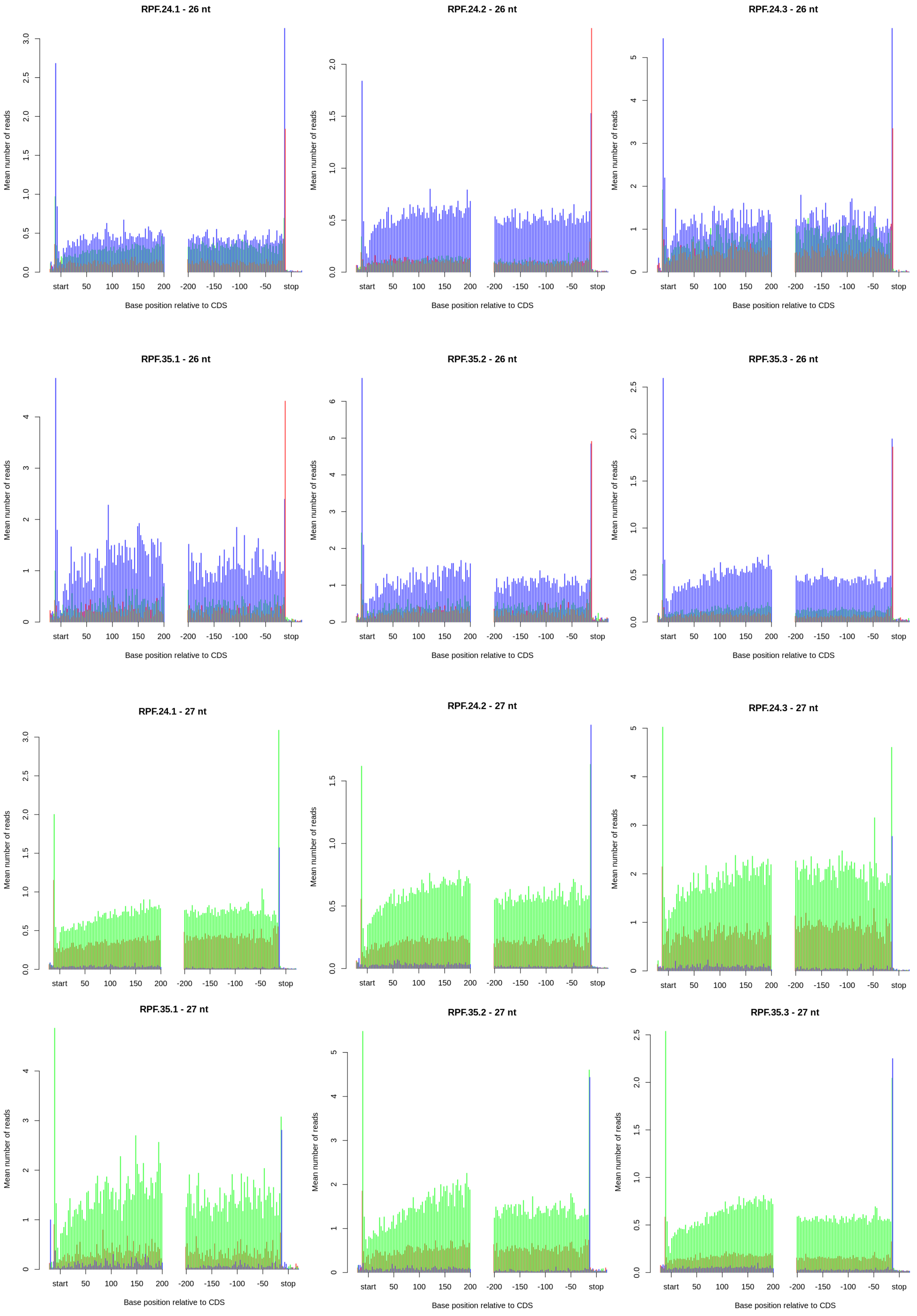
RPF.24.1 - 26 nt
RPF.24.2 - 26 nt
RPF.24.3 - 26 nt
RPF.35.1 - 26 nt
RPF.35.2 - 26 nt
RPF.35.3 - 26 nt
RPF.24.1 - 27 nt
RPF.24.2 - 27 nt
RPF.24.3 - 27 nt
RPF.35.1 - 27 nt
RPF.35.2 - 27 nt
RPF.35.3 - 27 nt
Mean number of reads
Base position relative to CDS
start
50
100
150
200
-200
-150
-100
-50
stop

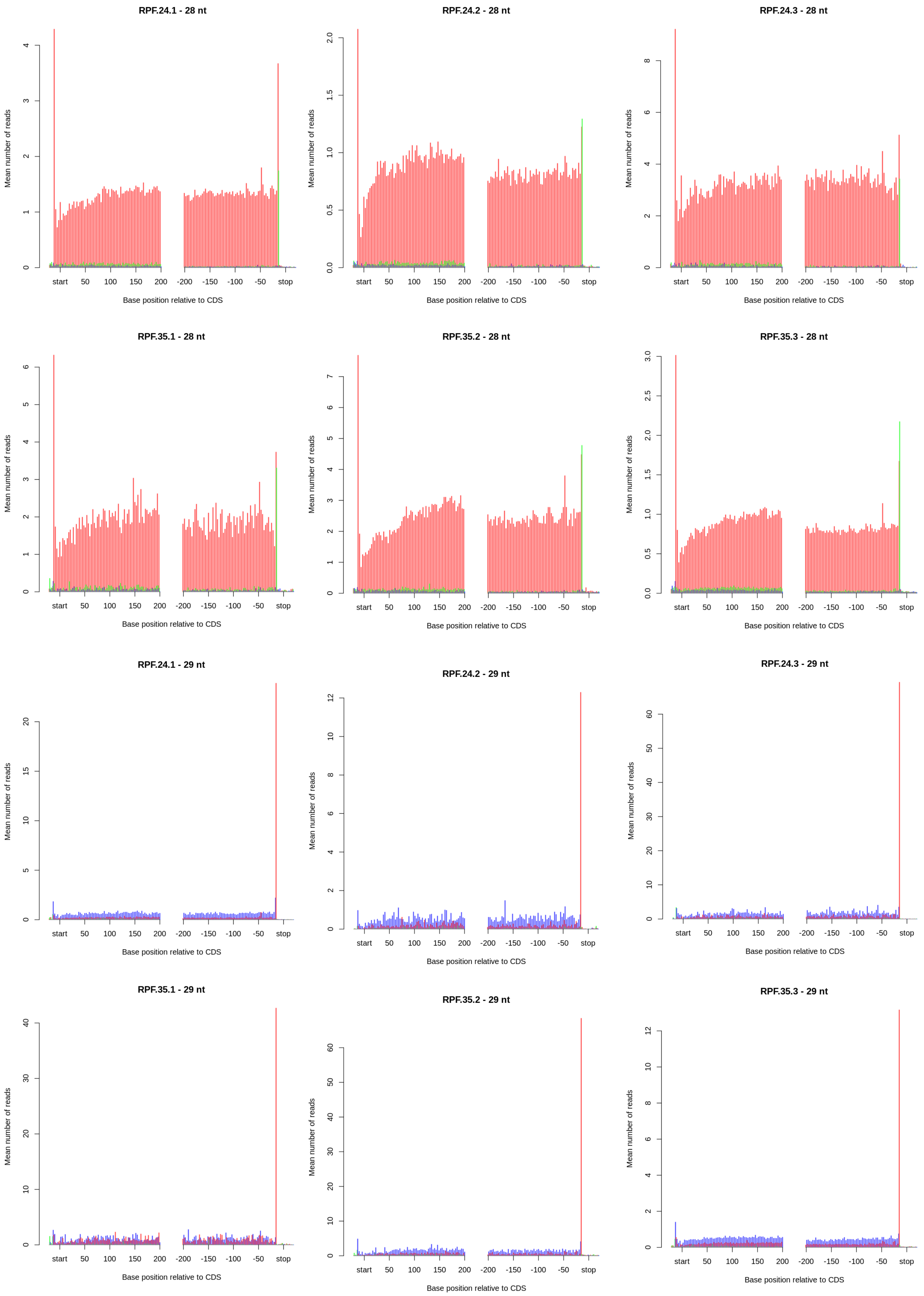

RPF.24.1 - 28 nt
RPF.24.2 - 28 nt
RPF.24.3 - 28 nt
RPF.35.1 - 28 nt
RPF.35.2 - 28 nt
RPF.35.3 - 28 nt
RPF.24.1 - 29 nt
RPF.24.2 - 29 nt
RPF.24.3 - 29 nt
RPF.35.1 - 29 nt
RPF.35.2 - 29 nt
RPF.35.3 - 29 nt
Mean number of reads
start 50 100 150 200 -200 -150 -100 -50 stop
Base position relative to CDS

### Figure S3

A

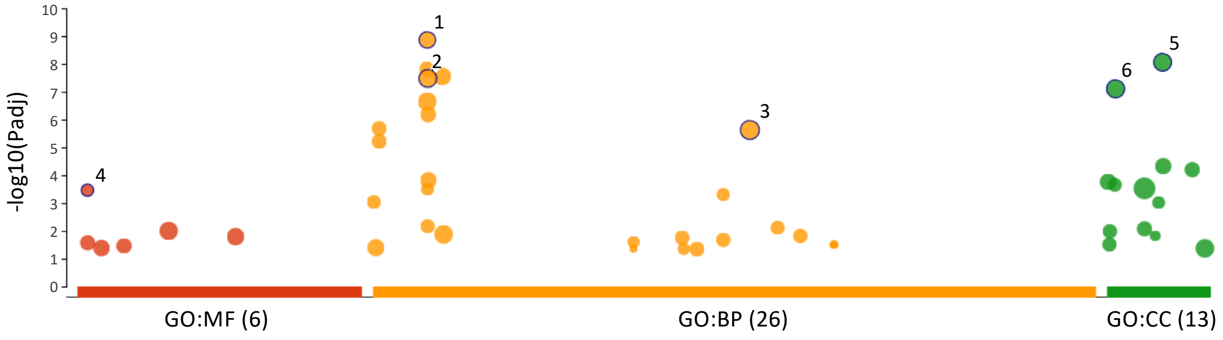


| ID | Term ID | Term Name | padj |
|---|---|---|---|
| 1 | GO:0030599 | DNA replication | $1.376\times10^{-9}$ |
| 2 | GO:0009860 | DNA repair | $3.330\times10^{-8}$ |
| 3 | GO:0009932 | chromosome organization | $2.360\times10^{-6}$ |
| 4 | GO:0016192 | DNA replication origin binding | $3.452\times10^{-4}$ |
| 5 | GO:0009856 | chromosomal part | $8.721\times10^{-8}$ |
| 6 | GO:0048588 | chromosome | $7.846\times10^{-8}$ |

B

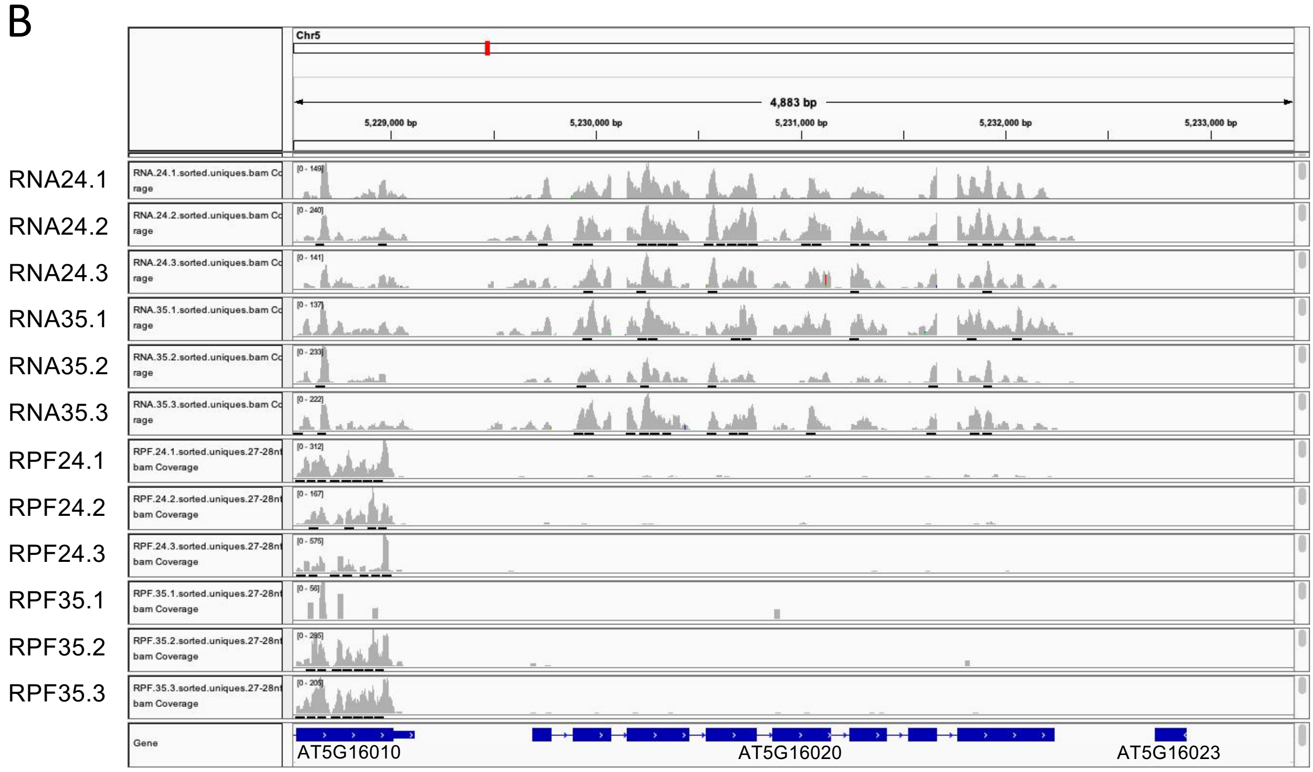
